# Exploring the mobilome and resistome of *Enterococcus faecium* in a One Health context across two continents

**DOI:** 10.1101/2022.04.11.487771

**Authors:** Haley Sanderson, Kristen L. Gray, Alexander Manuele, Finlay Maguire, Amjad Khan, Chaoyue Liu, Chandana N. Rudrappa, John H. E. Nash, James Robertson, Kyrylo Bessonov, Martins Oloni, Brian P. Alcock, Amogelang R. Raphenya, Tim A. McAllister, Sharon J. Peacock, Kathy E. Raven, Theodore Gouliouris, Andrew G. McArthur, Fiona S. L. Brinkman, Ryan C. Fink, Rahat Zaheer, Robert G. Beiko

## Abstract

*Enterococcus faecium* is a ubiquitous opportunistic pathogen that is exhibiting increasing levels of antimicrobial resistance (AMR). Many of the genes that confer resistance and pathogenic functions are localized on mobile genetic elements (MGEs), which facilitate their transfer between lineages. Here, features including resistance determinants, virulence factors, and MGEs were profiled in a set of 1273 *E. faecium* genomes from two disparate geographic locations (in the UK and Canada) from a range of agricultural, clinical, and associated habitats. Neither lineages of *E. faecium* nor MGEs are constrained by geographic proximity, but our results show evidence of a strong association of many profiled genes and MGEs with habitat. Many features were associated with a group of clinical and municipal wastewater genomes that are likely forming a new human-associated ecotype. The evolutionary dynamics of *E. faecium* make it a highly versatile emerging pathogen, and its ability to acquire, transmit, and lose features presents a high risk for the emergence of new pathogenic variants and novel resistance combinations. This study provides a workflow for MGE-centric surveillance of AMR in *Enterococcus* that can be adapted to other pathogens.

## 1 Introduction

*Enterococcus faecium* is a ubiquitous, gram-positive, facultative anaerobic microorganism often isolated from a variety of natural environments including soil and water, and host-associated environments including the intestinal tract of humans and animals [1–3]. The presence of *E. faecium* in the intestinal tract of healthy subjects led to the belief that this microbe was an innocuous commensal, with an occasional role in opportunistic infections [4]. However, following a 1986 outbreak of vancomycin-resistant strains of *Enterococcus faecalis* and *E. faecium* in a London, UK, hospital [5], it became clear that this bacterium could cause grave illness in humans and, readily acquired antimicrobial resistance (AMR) genes through lateral gene transfer (LGT). Enterococci have the ability to share genes within an extended pool of mobile genetic elements (MGEs) [6], allowing them to serve as hubs for the transmission of AMR determinants to both gram-positive and gram-negative species [7]. Antimicrobial-resistant enterococci are now a leading cause of hospital-acquired bloodstream and urinary tract infections [4]. According to the World Health Organization, *E. faecium* is a member of a group of nosocomial pathogens called “ESKAPE” [8] that have been given priority status on the list of pathogens for which new antimicrobials are urgently needed [9]. Up to the early 1990s, most nosocomial enterococcal infections were caused by *E. faecalis*, while *E. faecium* was the causative agent of only about 10% of cases [10]. Over the past two decades, *E. faecium* infections have been constantly on the rise in the United States [11–13] and in Europe [14–18]. In 2021, Dadashi *et al*. reported that the global prevalence of *E. faecium* among enterococci isolates from clinical infections was 40.6%, with 43.6% in Asia, 38.0% in Europe, and 36.8% in America [19].

Lebreton *et al*. [20, 21] described how *E. faecalis* and *E. faecium* have emerged independently through separate events of LGT driven largely by MGEs. Specifically, in *E. faecium*, there is a deep split (about 3,000 years ago) between strains commonly present in the microbiota of non-human animals (Clade A), which are the ancestors of most of the current clinical, and human-adapted commensal strains (Clade B). This split coincides with the loss of many genes related to the catabolism of dietary carbohydrates from Clade A strains and the MGE-mediated acquisition of genes encoding amino-carbohydrates typically involved in the glycocalyx formation during colonization of intestinal epithelial cells [20, 21]. The authors of these studies hypothesize that this difference in tropism is a reflection of the preferred habitats between these two clades, with Clade B mostly community-associated and Clade A mainly with hospital-associated enterococci [22]. In studies from the United Kingdom (UK) by Gouliouris *et al*. [23] and Alberta, Canada (AB) by Zaheer *et al*. [24], isolates from agricultural environments clearly separated from clinical ones constituting two distinct clades, supporting the hypothesis that they are specialized to distinct ecological niches. This adaptation also reflects the nature of antimicrobials, heavy metals, and other selective pressures present in each niche. Gouliouris *et al*. also included a clear split between Clade A subclades, A1 and A2, although we do not investigate these subclades in this study [23].

*E. faecium* is extremely apt at acquiring genes carried by MGEs including plasmids, genomic islands (GIs), and prophages. In fact, the genome plasticity that renders this microorganism a formidable public-health threat relies mainly on a large number of multifunctional accessory genes that can be laterally transferred between distantly related strains [6]. Plasmids are generally considered the main AMR gene-carrying MGEs in enterococci [25, 26]. Arredondo-Alonso *et al*. proposed that plasmids could be used to ascertain the niche specificity of *E. faecium* [27]. However, AMR, heavy metal resistance (HMR), and virulence factor (VF) genes have also been detected in GIs [28] and prophages [29].

Understanding the relative importance of habitat and geography in shaping the genome and corresponding resistance of *E. faecium* is vital for guiding antimicrobial use and AMR mitigation strategies. A “One Health” perspective that considers the emergence, dissemination, and transmission of resistance among human, agricultural, and environmental isolates is necessary to implement effective AMR surveillance and interventions. Existing surveillance systems rely heavily on phenotypic data and will benefit from whole-genome sequencing and analysis. The tools employed in this genomic study can connect phylogenetic, habitat, and geographic data to the prevalence of the mobilome and associated resistance and virulence genes, improving our knowledge of AMR dynamics in *Enterococcus* and other pathogens. Here we examine a combined set of 1273 *E. faecium* genomes from the United Kingdom and Alberta, Canada isolated from multiple habitats in order to determine the relationship between habitat, geography, and the occurrence and distribution of the mobilome and resistome of this opportunistic pathogen.

## 2 Methods

### 2.1 Genome Assembly and Classification

The dataset for this study encompassed 1766 genomes: 334 *E. faecium* genomes from Alberta, Canada [24] and 1432 from the United Kingdom [23]. These genomes originated from isolates collected from five different sources: Clinical (CLIN), Agriculture (AGRI), Municipal Wastewater (WW-MUN), Agricultural Wastewater (WW-AGR), and Natural Water sources (NWS). FASTQ files for the AB genomes were retrieved from the Sequence Read Archive (SRA) of the National Center for Biotechnology Information (BioProject PRJNA604849), and UK genomes from the European Nucleotide Archive (Table S1). The quality of the FASTQ files was determined using FastQC v0.11.8 [30]; reads were trimmed using fastp v0.23.2 and assembled with Unicycler v0.4.8 [31] using default parameters. The quality of the assemblies was assessed using Quast v5.0.2 [32] (Figure 1A). An NG50 cutoff of 30,000 bp was used to remove low-quality genomes from subsequent analysis.

**Fig. 1.**
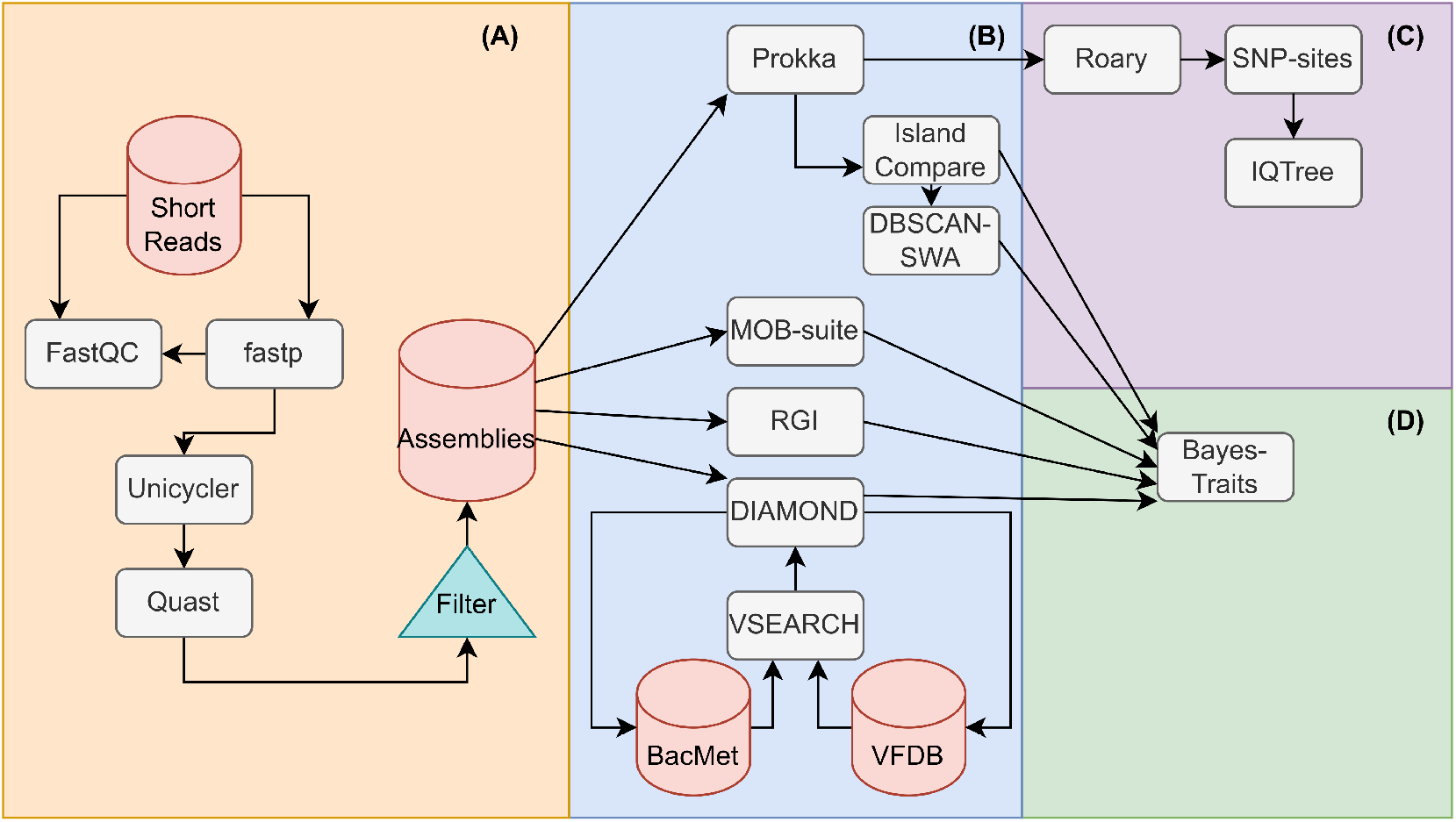
Overview of data processing workflow. (A) Short reads were trimmed using fastp, with quality checks by FastQC before and after trimming. The reads were then assembled using Unicycler and quality checked using QUAST. (B) The quality-checked assemblies were annotated with Prokka, MOB-suite, RGI, and DIAMOND. DIAMOND annotation was performed using the VFDB and BacMet databases. The Prokka annotations were passed to IslandCompare to infer probable GIs. The outputs from IslandCompare were passed to DBSCAN-SWA to infer probable phages. (C) Annotated contigs from Prokka were passed to Roary for pangenome calculation and core-genome alignment. The core genome alignment’s static sites were removed using SNP-sites and the resultant alignment was passed to IQ-Tree to calculate a maximum likelihood phylogenetic tree. (D) All annotated target genes and MGEs were tabulated and passed to BayesTraits for co-evolutionary analysis, performing hypothesis testing for correlated evolution between pairs of features.

### 2.2 Pangenome and Generation and Visualization of Core-Genome Phylogenies

We constructed a core-genome phylogenetic tree for all of the *E. faecium* genomes. To do so, genomes were annotated using Prokka version 1.14.6 [33] followed by Roary v3.13.0 to construct a core-genome alignment [34]. As a reference, the genome of *E. faecium* DO ASM17439v2 was included and *E. hirae* ATCC9790 ASM27140v2 was used as an outgroup genome in the alignment. Using SNP-sites v2.5.1 a single-nucleotide polymorphism (SNP) alignment was produced and unambiguous nucleotide frequencies were counted [35]. The resultant SNP alignment and core-genome base frequencies were then used to generate a maximum-likelihood phylogenetic tree using IQTree v.2.1.4-beta [36] with the general time reversible model with invariable site plus discrete gamma model (Figure 1C). One thousand ultrafast bootstrap replicates and 1000 Shimodaira Hasegawa-like approximate likelihood ratio tests (SH-aLRT) were performed [37]. The phylogeny was then visualized using GrapeTree v.1.5.0 [38].

Genomes were assigned to “Clade A” and “Clade B” based on *groEL* gene sequences as described in Hung *et al*. [39]. Briefly, *groEL* sequences were extracted and sequences corresponding to “Clade A” strains *E. faecium* strain V68, accession MH109129 and “Clade B” *E. faecium* strain 81, accession MH109127 were added as references. Sequences were aligned using MAFFT v7.490 with default parameters and a maximum-likelihood tree was created with IQTree v2.1.4 using as a model unequal purine/pyrimidine rates with empirical base frequencies and a proportion of invariable sites (TN+F+I). Our assignment of the genomes to the two categories was then guided by the topology of this tree. Habitats, countries of origin, and “Clades” were all mapped onto the core genes-based reference tree. “Clade B” was paraphyletic in the reconstructed tree, as a consequence we refer to these two categories as “Type A” for “Clade A” and “Type B” for “Clade B”.

### 2.3 Prediction of Resistance Genes and Virulence Factors

The assembled contigs were used to detect AMR genes, HMR genes, and VFs (collectively referred to as target genes) using specific databases for each gene type. AMR genes were detected with the Resistance Gene Identifier (RGI) (v5.1.0) for the prediction of AMR genes based on homology and SNP models from the Comprehensive Antimicrobial Resistance Database (CARD) v3.1.0 [40]. RGI stratifies matches into three categories using a curated BLAST bitscore cutoff that reflects known variation within a gene. Matches that score below a designated, target-specific cutoff are assigned to the “Loose” category, while matches above this cutoff are assigned to the “Strict” category. Exact matches are assigned to the “Perfect” category. We limited our analysis to the Strict and Perfect matches. To identify HMR genes and VFs, open reading frames (ORFs) present in the assemblies were first annotated using Prodigal v2.6.3 [41]. Subsequently, homology search with an initial E-value threshold of 10^−20^ against two databases, VFDB [42] and BacMet v2.0 [43] was conducted using DIAMOND-BLASTX v0.9.36 [44]. Results were then filtered by percent identity (60%) and match coverage (60%). Clusters of highly similar genes (> 95%) were identified using vsearch v2.17.1 [45] (Figure 1B).

### 2.4 Prediction of Mobile Genetic Elements

To predict the presence of plasmids in our short-read assemblies, the genomes were analyzed with MOB-suite 3.0.0 [46] with the v2020-05-05 database. The MOB-suite pipeline scans input assemblies for contigs containing plasmid-related genes (e.g. relaxases and replicases) and repetitive regions, thereby identifying putative plasmid scaffolds. These scaffolds were compared against a database of mobility clusters (MOB-clusters) comprising pre-clustered reference plasmids. The putative plasmids were assigned to MOB-clusters by identifying the minimum Mash distance [47] to a reference plasmid. The output consisted of a single contig sequence per MOB-cluster and an annotation of their host-range predictions, mobility predictions, and assignment to a replicase (*rep*) gene cluster. Contigs larger than 1 kb were examined for the presence of GIs with IslandCompare [48] using the reference genome *E. faecium* DO ASM17439v2. IslandCompare uses the reference genome to generate an alignment-based concatenated genome from each submitted draft genome. GIs are predicted by two underlying tools, IslandPath-DIMOB [49] and Sigi-HMM [50] that identify regions of the genome with anomalous dinucleotide bias (and at least one mobility gene) and differential codon usage, respectively. IslandCompare also incorporates additional functionality for ensuring the consistency of GI predictions across genomes in multi-genome datasets and clusters the predicted GIs. Following analysis, any GI predictions that corresponded to the region of the genome that could not be aligned to the reference genome were excluded. For GIs present in a relatively large proportion of genomes (> 10%), a manual assessment was performed for genes annotated in those GIs. One GI that consisted mainly of genes involved in replication was present in nearly all genomes (1208/1273) and excluded from the analysis.

Concatenated genome files generated by IslandCompare were also used for prophage prediction via DBSCAN-SWA [51]. DBSCAN-SWA combines density-based spatial clustering of applications with noise (DBSCAN) and a sliding window algorithm (SWA) for prophage detection (Figure 1B).

The taxonomic distribution of predicted genes and MGEs was assessed through a homology search. A reference database of predicted proteins was constructed from 20,100 complete bacterial genomes downloaded from RefSeq on December 24, 2021. DNA sequences of plasmid-associated contigs, GIs, and prophages were compared to this database using DIAMOND-BLASTX version 2.0.13 with a maximum e-value of 10^−50^, 90% percent identity or greater, and minimum subject coverage of 90%. Only matches with a score of 95% or greater relative to the best match were retained. Final filtering of results used a minimum percentage identity threshold of 99%. The taxonomic distribution of matches was extracted from the resulting set of hits.

Sets of target genes that mapped to a given MGE were considered to be co-localized to that MGE. Co-localizations between predicted AMR, VF, and HMR genes were identified using python and summarized by gene cluster. Genes that did not localize to any MGE were treated as chromosomal.

### 2.5 Analysis of Feature Abundance

We tested the hypothesis that *E. faecium* isolates from different sampling environments have differential abundances of AMR and HMR determinants, VFs, and MGEs (hereafter referred to collectively as “features”). A three-way factorial ANOVA was performed to determine the extent that features were associated with habitat, geographic location, type, and the interactions of these categories.

To perform ANOVA for each feature, the mean number of unique features of each type per genome was calculated. Genomes originating from NWS and WW-AGR isolates were available only in the AB isolates so these genomes were omitted. This resulted in 1203 genomes divided over 12 treatment groups (3 habitats, 2 geographic locations, 2 types). To account for the unequal number of isolates in each treatment group, we used unweighted marginal mean sum of squares (SS), or type III SS, to calculate our ANOVA statistics [52]. To investigate feature frequency variance in the omitted environments, a two-way ANOVA was performed using the 303 AB genomes with 10 treatment groups (5 habitats and 2 types). Where categories were found to be significant at *α* = 0.05, pairwise Tukey’s HSD post-hoc significance testing was performed on within-category group means (Table S2).

### 2.6 Coevolutionary Associations of Target Genes and Mobile Genetic Elements

We investigated the correlation across and within features using phylogenetic profiles. A phylogenetically informed maximum-likelihood approach was used to predict pairs of features with coordinated patterns of gain and loss, using the BayesTraits version 3.0 [53]. To characterize the associations of predicted genes and MGEs across the tree, BayesTraits constructs two continuous-time Markov models for each gene/gene, gene/MGE, and MGE/MGE pair based on their patterns of presence and absence: one model expresses the likelihood that the pair evolves in a correlated way, and the other the likelihood that they are gained and lost independently [54, 55]. The ratio of these two likelihoods was used to generate a p-value that reflects the statistical significance of their association. Only pairs in which both features occurred in at least 3 genomes were considered. Because directionality of the association is not determined by BayesTraits, we infer this by referring to the distribution of the genes across the phylogenetic tree, habitat, clade and geography using the presence and absence of the features.

The likelihood ratios and p-values corresponding to the gene-gene and gene-environment relationships were represented as network diagrams using Cytoscape (v3.8.2).

## 3 Results

### 3.1 Genome Assembly and Distribution

Out of 1766 sequenced genome initially selected for this study, 1273 genome assemblies (i.e., 303 from AB and 970 from UK) that passed quality-control measures (NG50 >= 30,000) were selected for further analysis. The NG50 values of accepted genomes ranged from 30,088 to 467,170 bp, with a mean of 148 contigs (range: 18 to 304). Assembly sizes varied from 2,373,576 bp to 3,301,308 bp with a mean value of 2,830,264 bp (Figure 2A) and with a mean GC content of 37.8% (range from 37.25% to 38.49%). The pangenome of the 1273 *E. faecium* genomes consisted of 26,246 genes (Table S3), including 1101 (4.2%) present in 99-100% of the genomes (the core genome); 212 (0.8%) in 95-99% of genomes (the “soft core” genome); 2207 genes (8.4%) in the “shell” genome (15-95% of genomes), and 22,726 genes (86.6%) in the “cloud” (less than 15% of genomes). A total of 382 genes belonging to the AMR (n = 82), HMR (n = 32), and/or VF (n = 268) classes were identified at different frequencies throughout the analyzed genomes (Figure 2B-H).

**Fig. 2.**
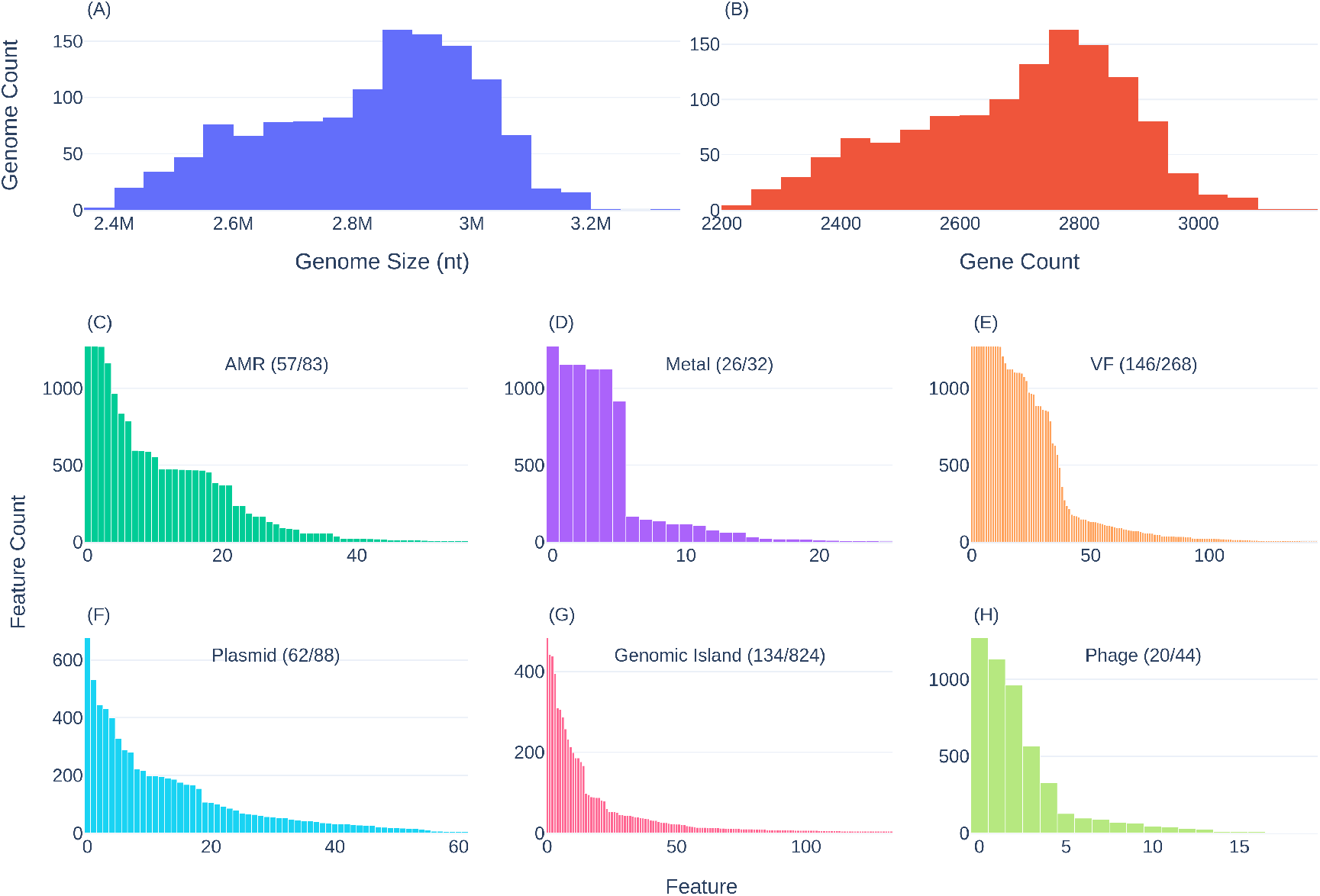
Size and count distributions of genomic features. First row: Genome size (A) and Pan-genome distribution predicted by Roary (B). Second row: frequency distribution of AMR genes (C), HMR genes (D), and VFs (E). Third row: frequency distribution of plasmids (F), GIs (G), and phages (H). Multiple occurrences of a feature in a given genome were counted only once. For clarity, only features detected in at least five genomes were plotted. Plot annotations indicate the number of features plotted and the number of total features detected.

Our analysis of the predicted mobilome identified 1131 sets of MGEs, all of which were part of the accessory genome except for *Streptococcus* phages predicted by DBSCAN-SWA in 1272 genomes. MOB-suite assigned plasmid-associated contigs to 263 different clusters. Of these, 88 were assigned to reference plasmid clusters and 175 were categorized as novel with 144 (83.2%) of these associated with UK isolates. Given that many of these novel clusters may be misclassified chromosomal fragments, we did not include these in our subsequent analysis. The 7805 GIs were predicted to constitute 824 groups of GIs with only 15 GI clusters present in more than 10% of the isolates, 477 of them being unique to single genomes.

### 3.2 Distribution of the Resistome, Virulence Factors, and MGEs by Type Assignment, Geographical Origin, and Habitat

The factorial three-way ANOVA compared the main effects of habitat, geography, and type as well as their interaction on the frequency of target features (Table 1). For all features, marginal effects at a threshold of *α* = 0.05 were observed for at least one of the categories (geography, habitat, and type). However, in each case at least one significant interaction effect was also observed which suggests conditionally dependent patterns of association and the need to interpret main effects with caution. Habitat showed the strongest marginal effects for all features except HMR, with *p* < 10^−7^ in all other cases. Geography was associated with HMR, VFs, and plasmid clusters, but post-hoc significance testing strongly suggested that the HMR relationship (*p* = 4.16 × 10^−5^) is an artefact of interaction effects driven by Type A agricultural isolates from the UK (Table S2B). AMR and phage showed significant associations with the Type A / Type B split, although the latter (*p* = 4.69 × 10^−2^) was not supported by post-hoc testing. In the two-way ANOVA using Alberta genomes, results were consistent with the three-way ANOVA with the exception of HMR determinant frequencies, which were found to be significantly affected by habitat at p<0.05 (Table 2). Inspection of the HMR frequency post-hoc significance tests indicate that this effect is likely caused by habitat × type interactions, with NW and CLIN isolates showing significant differences between Type A and Type B HMR determinant frequencies (Table S2B).

**Table 1.**
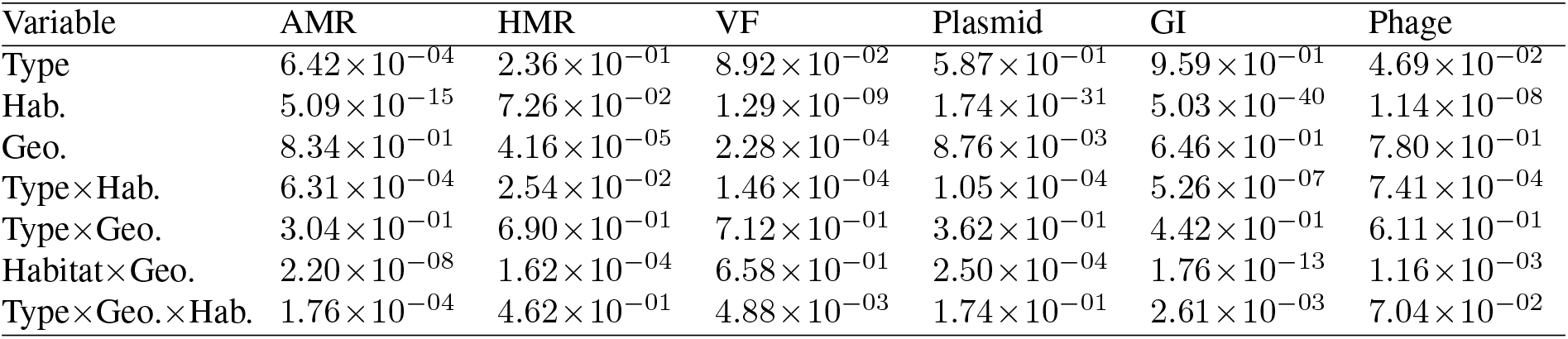
Three-way ANOVA results for 1200 genomes in habitats that were sampled in both AB and UK. Type III error correction was used to account for imbalanced classes. Columns indicate p-values testing differences of mean unique features per genome. Factor1 × Factor2 indicates interaction effects among categories.

| Variable | AMR | HMR | VF | Plasmid | GI | Phage |
| --- | --- | --- | --- | --- | --- | --- |
| Type | $6.42 \times 10^{-04}$ | $2.36 \times 10^{-01}$ | $8.92 \times 10^{-02}$ | $5.87 \times 10^{-01}$ | $9.59 \times 10^{-01}$ | $4.69 \times 10^{-02}$ |
| Hab. | $5.09 \times 10^{-15}$ | $7.26 \times 10^{-02}$ | $1.29 \times 10^{-09}$ | $1.74 \times 10^{-31}$ | $5.03 \times 10^{-40}$ | $1.14 \times 10^{-08}$ |
| Geo. | $8.34 \times 10^{-01}$ | $4.16 \times 10^{-05}$ | $2.28 \times 10^{-04}$ | $8.76 \times 10^{-03}$ | $6.46 \times 10^{-01}$ | $7.80 \times 10^{-01}$ |
| Type × Hab. | $6.31 \times 10^{-04}$ | $2.54 \times 10^{-02}$ | $1.46 \times 10^{-04}$ | $1.05 \times 10^{-04}$ | $5.26 \times 10^{-07}$ | $7.41 \times 10^{-04}$ |
| Type × Geo. | $3.04 \times 10^{-01}$ | $6.90 \times 10^{-01}$ | $7.12 \times 10^{-01}$ | $3.62 \times 10^{-01}$ | $4.42 \times 10^{-01}$ | $6.11 \times 10^{-01}$ |
| Habitat × Geo. | $2.20 \times 10^{-08}$ | $1.62 \times 10^{-04}$ | $6.58 \times 10^{-01}$ | $2.50 \times 10^{-04}$ | $1.76 \times 10^{-13}$ | $1.16 \times 10^{-03}$ |
| Type × Geo. × Hab. | $1.76 \times 10^{-04}$ | $4.62 \times 10^{-01}$ | $4.88 \times 10^{-03}$ | $1.74 \times 10^{-01}$ | $2.61 \times 10^{-03}$ | $7.04 \times 10^{-02}$ |

**Table 2.**
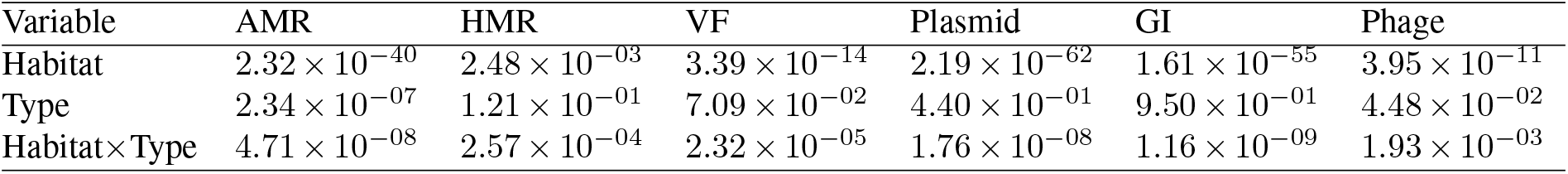
Two-way ANOVA for 303 AB genome assemblies, performed separately to account for the WW-AGR and NWS habitats exclusive to AB. Type III error correction used to account for imbalanced classes. Columns indicate significance values testing differences of mean unique features per genome. “Habitat × Type” indicates the interaction effect of habitat and type categories.

A total of 19,599 AMR genes was predicted in the 1273 isolates: removal of duplicates yielded 16,898 genes with a mean occurrence of 13.27 per genome. We observed a large discrepancy between types A (14.3±5.3 predicted AMR genes per genome) and B (5.9±1.6 AMR genes per genome) (Table S4; Figure 3). Plasmids and GIs showed discrepancies as well, with 6.2±3.0 mean occurrences of distinct plasmid clusters in Type A and 2.0±1.3 in Type B. GIs were found 7802 times, with 6.5±2.8 of mean occurrences in Type A and 3.7±1.6 in Type B. Conversely, HMRs, VFs, and phages showed similar distributions between types even when genomes were partitioned by geography and habitat.

**Fig. 3.**
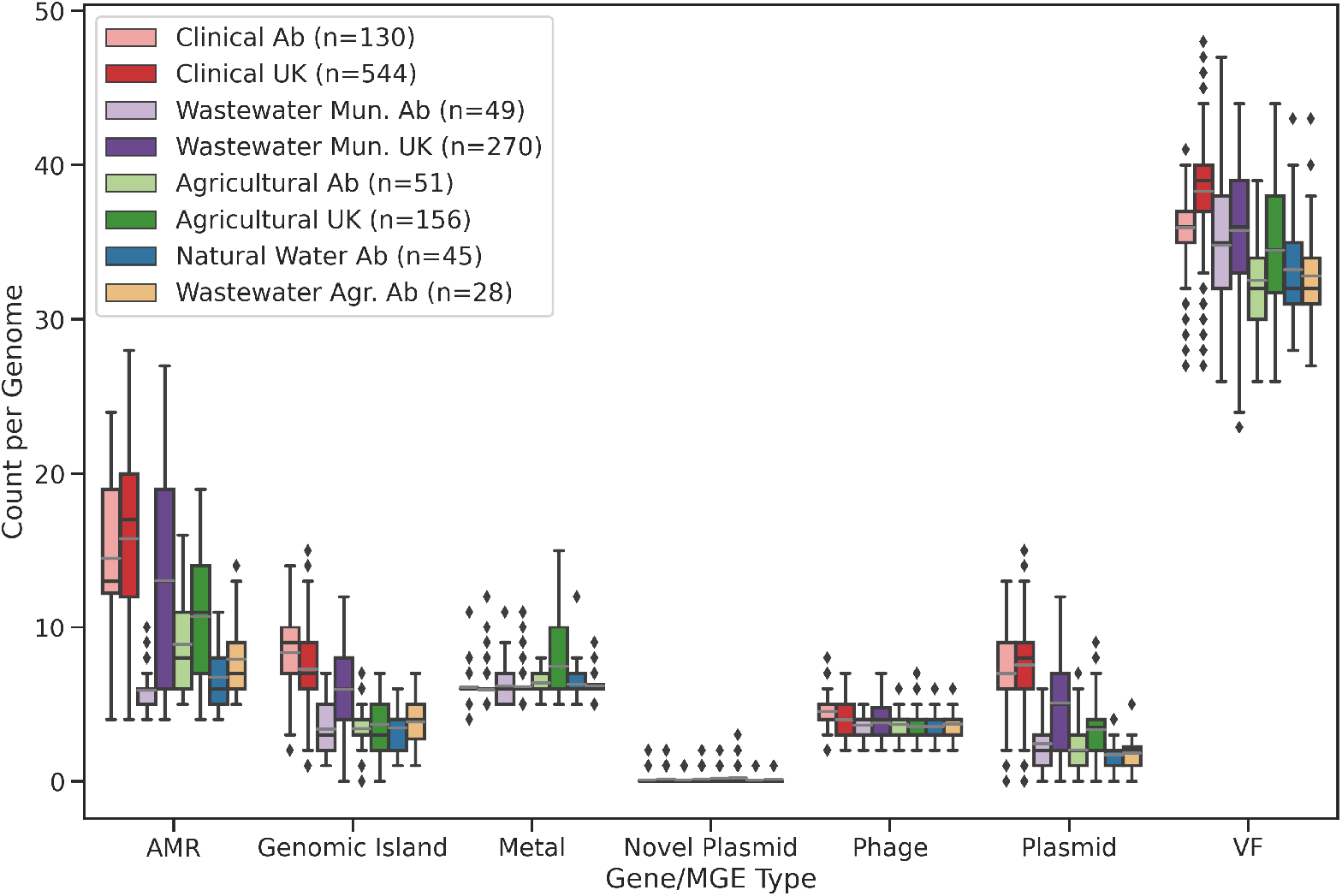
Abundance of features by habitat type and geographic location. “AB” indicates genomes sampled from Alberta, Canada. “UK” indicates genomes sampled from the United Kingdom. Counts indicate the number of unique features of a given category found per genome. Bars indicate quartiles. Points/diamonds are considered to be outliers if they fall outside 1.5 × the interquartile range. Grey bars indicate mean values.

For AMR genes, plasmids, and GIs, the difference in distribution between types A and B was still observed after considering the geographical origin of the samples (Table S4). However, we detected a large variation in the distribution of the AMR genes, plasmids, and GIs between UK and AB samples isolated from municipal wastewater. Specifically, the UK Type A genomes had 14.3±6.4 AMR genes, 5.5±3.0 plasmids, and 6.3±2.3 GIs compared to the AB Type A genomes with 6.44±1.9 AMR genes, 2.7±1.9 plasmids, and 3.2±1.5 GIs (Table S4). Differences in the distribution of AMR genes, plasmids, and GIs between types A and B were also detected across habitats. Specifically, Type A CLIN samples from UK and AB had the highest mean of AMR genes per genome of all the isolates (15.9±4.5 and 15.0±4.0 respectively) with corresponding variation in the relative distributions of plasmids and GIs.

### 3.3 Phylogenetic associations of target genes and MGEs

The phylogenetic tree inferred from the 1273 genomes separates isolates by type (Figure 4). Over half of Type A was comprised of almost entirely CLIN and WW-MUN isolates (mainly consisting of UK isolates), but two additional subclades possessed isolates from all habitats. Type B encompassed isolates collected from both countries and all five habitats, with environmental and geographical categories constituting monophyletic groups.

**Fig. 4.**
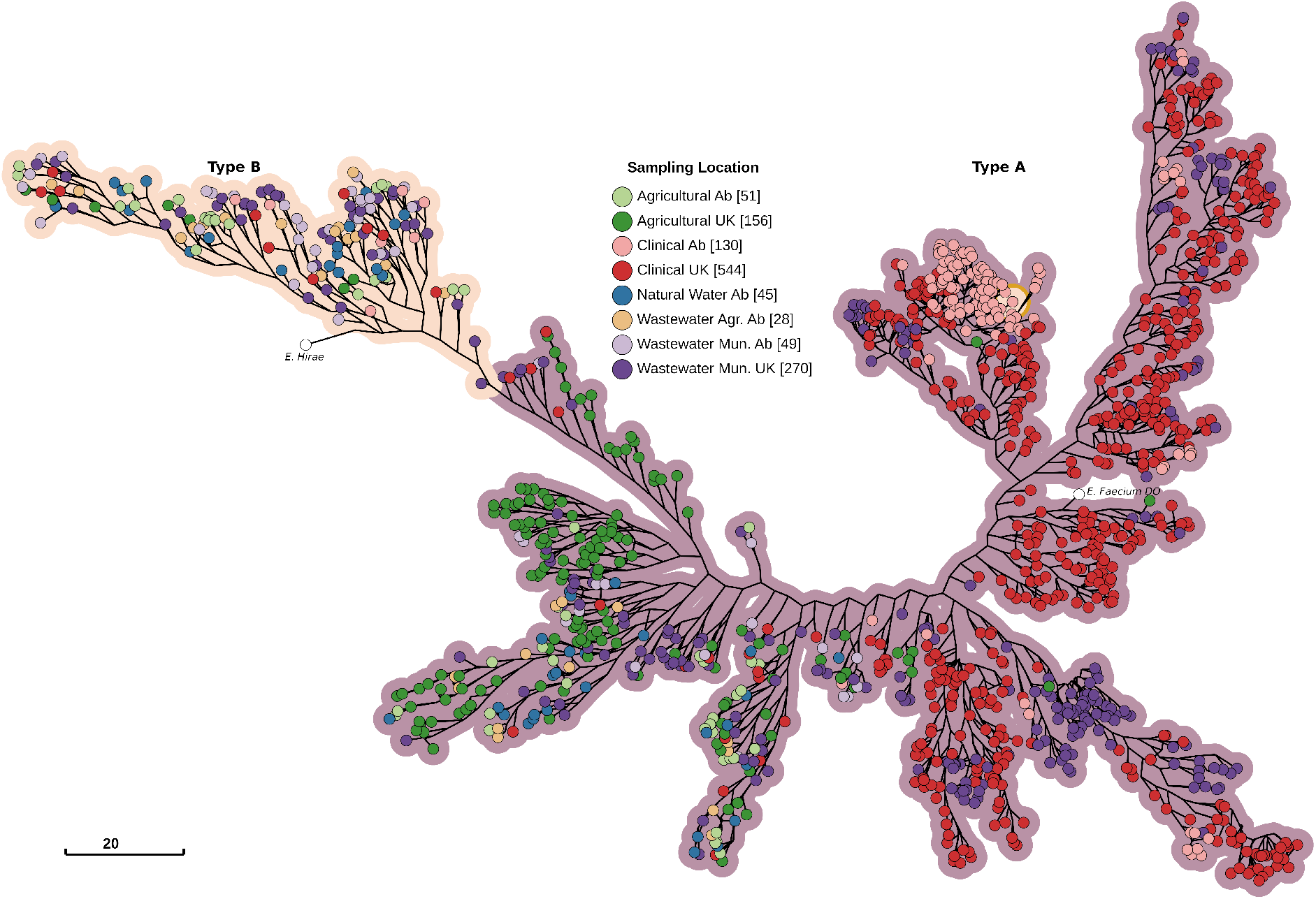
Maximum-likelihood core-genome phylogenetic tree of 1273 *E. faecium* genomes with *E. hirae* ATCC9790 as the outgroup, and *E. faecium* DO ASM17439v2 reference genome. The tree was constructed with 1,854,991 nucleotide sites, 79,440 of which were parsimony informative, using the general time reversible substitution model with invariant sites and four Gamma rate categories. Branch lengths are log-transformed and scaled down to 13% length for improved readability. Nodes are colored by sampling location, with hue indicating habitat and saturation indicating geography.

Most features were irregularly distributed across the phylogenetic tree (Figures S1A-F). Relatively few AMR genes were present in Type B compared to Type A, particularly Type A CLIN. Only *eatAv*, a variant of *eatA* that confers resistance to multiple antibiotics, was present in the majority of non-clinical subtrees and absent from most CLIN and WW-MUN isolates. Both the *vanA* and *vanB* operons were restricted to the CLIN / WW-MUN subtree. Two sets of HMR genes showed strong negative associations, largely mapping onto the Type A / Type B division; the corresponding genes had the same names (*chtR, ruvB, copB*, and *chtS*). These clusters may have been divided because of sequence dissimilarity rather than functional differences. Although some VFs were preferentially associated with Type A or Type B isolates, very few were exclusively confined to one or the other. Some plasmids and GIs were over-represented in the CLIN / WW-MUN Type A, and less frequently associated with the non-clinical isolates and Type B.

Significant (*p*-values < 0.01) associations were observed for AMR features with geography (18.0%), type (12.5%), and habitat (11.5%; *p* < 10^−10^) (Figure S2A). Overall, the strongest positive and negative associations were seen for the CLIN and AGRI habitats, respectively. The *vanA* genes and genes conferring resistance to aminoglycosides, macrolides, tetracycline, trimethoprim, streptothricin all met this threshold (Table S4). As an example, *vanA* was prevalent in CLIN genomes but almost never present in AGRI genomes. VFs exhibited associations with habitat, geography and type (Figure S2A, S2C). HMR genes associated most strongly with geographic origin than habitat or type (Figure S2B).

### 3.4 Physical location of the resistome genes and virulence factors in the genome

We next examined the localization of the 72,786 predicted AMR, HMR, and VF genes in the 1273 genomes. A total of 18,518 (25.4%) predicted genes mapped to one or more MGEs, with 20.6% mapping to plasmids, 5.1% to GIs, and 2.2% to prophages. There were a total of 102 MGEs with colocalized AMR and VF genes and a total of 7 MGEs with both AMR and HMR genes (Table S5).

The dominant plasmid clusters AB369, AC731, and AB173 were identified in 400, 678, and 187 genomes, respectively. Common AMR genes in these plasmids included the *vanA* operon, *sat-4, ermB*, and the aminoglycoside resistance genes *aac(6’)-Ie-aph(2”)-Ia, aad(6*), and *aph(3’)-IIIa*. Six VFs associated with the PilA pilus structure were also frequently found on these plasmids. However, the relative numbers of these genes differed among predicted plasmid clusters, with some containing solely AMR or VF genes. AB756, the fourth most-abundant plasmid cluster, also had substantial numbers of the three aminoglycoside resistance genes mentioned above, *ermB*, and *sat-4*, along with *lsaE* and the tetracycline resistance genes *tetL, tetM*, and *tet(W/N/W)*. AH273, the seventh most-abundant plasmid cluster, contained the HMR genes encoding the UDP-glucose 4-epimerase *galE* and the copper-translocating ATPase *copA*.

The most common GIs mainly housed tetracycline- and vancomycin-resistance genes. GI 14 (identified in 186 genomes) was associated with *dfrG, tet(W/N/W)*, and tetM; GI 8 (identified in 310 genomes) with *dfrF;* and GI 34 (identified in 51 genomes) with the *vanB* suite of genes. GI 69 was found less frequently than the predominant GIs, being present in 18 genomes and containing *ant(9)-Ia, efrA*, and *ermA*. Other predicted GIs had aminoglycoside-resistance genes, *ermB* (a macrolide resistance gene), and *sat-4*. Predicted VFs in GIs included *bsh* (VFC36), a bile salt hydrolase, *ssaB* (VFC39), a Manganese/Zinc ABC transporter substrate-binding lipoprotein precursor, *fss3* (VFC42) a fibrinogen binding protein, *ecbA* (VFC84 and VFC86) a collagen binding protein, and multiple genes involved in capsule formation (*epsE* (VFC48), *gmd* (VCF51), *cps2K* (VCF52)).

Most prophage-associated genes mapped to either annotated *“Streptococcus* phage” (1366 / 1578) or *“Enterococcus* phage” (100 /1578). Genes that mapped to the predicted *Streptococcus* phages were similar to those observed in the plasmids and GIs, including those associated with aminoglycoside, erythromycin (*ermB*), streptothricin (sat-4), and tetracycline resistance. While some common VFs were unique (eg. *lap* (VFC14), an alcohol dehydrogenase involved in adhesion to the host cells), others were similar to those found to be localized to GIs including *bsh* (VFC22 and VFC36), *fss3* (VCF42), and *ecbA* (VFC84). Predicted *Enterococcus* phages had several instances of *dfrA42, bsh* (VFC22).

Many of the genes noted above showed biased associations with the corresponding MGEs. For example, over 93% of all *vanA* and tetracycline-resistance genes mapped to predicted plasmids, as were over 80% of the macrolide and streptothricin resistance genes *ermB* and *sat-4*, respectively. However, the gene-centric view also identified rare genes with strong biases including *catA8, lnuB, ermT*, and chloramphenicol acetyltransferases. Over 75% of *vanB* and *dfrF* genes were associated with GIs; other AMR genes with strong biases included *optrA* (65%) and *lnuG* (61.9%). The genes most strongly associated with prophages were the collagen-binding MSCRAMM gene (86.5% of genes), *tet(W/N/W)* and *tetM* (61.7% and 40.4% respectively) and *dfrG* (33.5%). However, all of these genes were also strongly associated with plasmids, GIs, or both. No prophage-specific genes were identified.

### 3.5 Phylogenetic Distribution of MGEs

We examined the predicted host-range distributions of all features via direct homology search and the host-range prediction feature of MOB-suite. MOB-suite predicted a total of 7232 putative plasmids which grouped into 88 unique MOB-clusters. A total of 4470 plasmid-associated contigs were predicted to be non-mobilizable, while 1782 were predicted to be mobilizable and 980 were predicted to be conjugative. The majority of plasmid-associated contigs (n=5652) were predicted to be specific to *Enterococcus*. Relatively few plasmid clusters had very narrow or very wide distributions outside of *Enterococcus:* a total of 358 clusters were associated with some other single genus, while 347 clusters were predicted to occur in phyla other than Firmicutes.

Although plasmid host range was often narrow, individual plasmid-associated genes were often strikingly similar (>99% identity over at least 90% of the subject sequence) to genes from other taxonomic groups, even at the phylum level. Of the contigs annotated as cluster AC731 across 678 genomes (36 mobilizable), 206 had at least one aminoglycoside-resistance gene with a high-stringency match outside of *Enterococcus*, frequently to *Staphylococcus*, *Streptococcus*, *Campylobacter*, and members of the family Enterobacteriaceae, while 112 matched at least one gene in the *vanA* group, often across multiple phyla. Not all plasmids showed evidence of recent LGT. All non-hypothetical genes in plasmid cluster AH273, including a range of metal-associated transporters, had no stringent matches outside of *Enterococcus*. All the annotated members of this plasmid cluster were predicted by MOB-suite to be non-mobilizable.

Most GIs were dominated by integrases and other signatures of MGEs, and poorly annotated genes with products that include general ABC transporters. The most common GI was found in 484 genomes; over 90% of these GIs had a suite of genes found in multiple phyla and included toxin-antitoxin and pilin genes, peptidases, and annotated ABC transporter permeases. GI 26, found in 88 genomes, had a very high incidence of multiphylum *tetM* genes. The *vanB* genes found in GI 34 were nearly identical to those in other Firmicutes such as *Staphylococcus*, and occasionally in members of Enterobacteriaceae such as *Klebsiella*. Similarly, prophage genes with stringent matches to groups outside *Enterococcus* were predominantly associated with mobility and included endonucleases, integrases, and transposases. However, over 500 genomes had genes annotated as *ermB*, with stringent matches to other phyla. A similar number of genomes had at least one aminoglycoside-resistance gene, the most common being *ant(6)-Ia*.. Other common genes involved in transcription included transcription factors and 500 instances of the σ^70^ subunit of RNA polymerase.

### 3.6 Distribution and Associations of Vancomycin-Resistance Genes

Both the *vanA* and *vanB* gene clusters showed a strong association with CLIN and WW-MUN but variable distribution and association with MGEs (Figure 5). The *vanA* gene clusters were found to be disproportionately associated with plasmids in both the AB and UK datasets (Table S5). Overall, 458/474 *vanA* genes were found to colocalize to and associate with several plasmids of which AB369 and AC731 were the most abundant. The *vanA* genes were primarily identified in CLIN (100/474 AB and 270/474 UK) and WW-MUN samples from the UK (103/474), with only 1 isolate from UK AGRI sources.

**Fig. 5.**
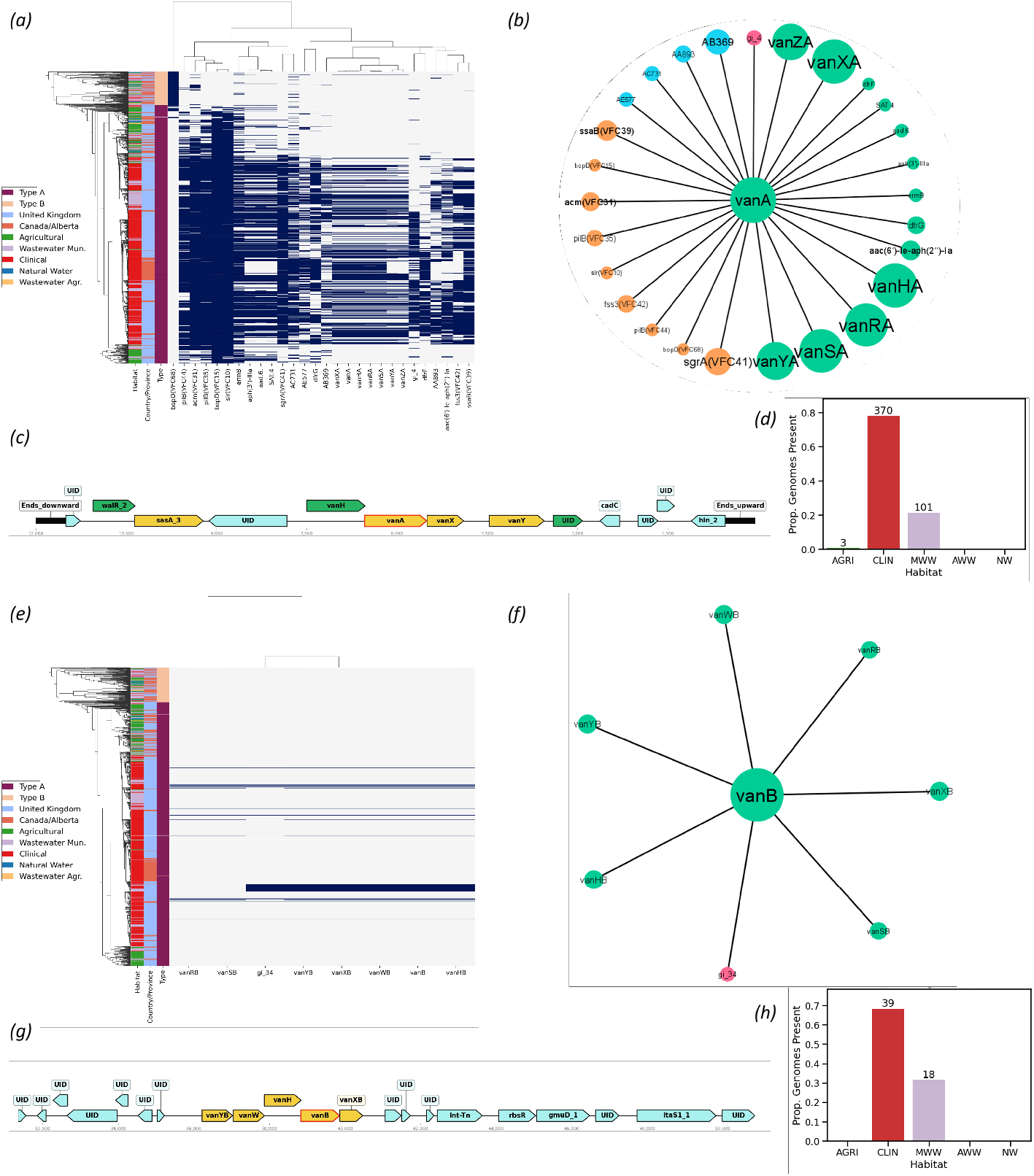
Statistical associations and physical localization of *vanA* (A-D) and *vanB* (E-H) genes. (A,E) Phylogenetic distribution of *van* genes and other features with an associated likelihood ratio ≥ 100. (B,F) Statistical association network of *vanA/vanB* genes with other features. Gene and MGE colours are consistent with those in Figure 2. (C,G) Example of gene order on an annotated plasmid (C) and GI (G) Green genes correspond to “Perfect” matches with reference genes in the CARD database, yellow genes are “Strict” hits. (D-H) Distribution of genes by habitat. Bar colors correspond to their habitats as per the legend in (A,E).

Conversely, all of the *vanB* gene clusters were found in UK genomes and, in 51/57 genomes, these genes colocalized to GIs (Figure 5). For the remaining 6/57 genomes, the *vanB* was in the unaligned portion of the genome that was not included in the GI analysis. All of the *vanB* genes were predicted to fall on a single GI cluster (except for one representative that has a large insertion in the middle of the GI) and contained a Tn916 transposase. The positive association of *vanB* to GI cluster 34 was also supported (*p* < 10^−16^). The majority (39/57) of *vanB* genes were in CLIN isolates with WW-MUN isolates composing the remaining 18/57 instances.

### 3.7 Other Notable AMR Gene Classes

In addition to *msrC* (Figure 6), a species-specific gene of *E. faecium* that confers low-level intrinsic resistance to macrolide and streptogramin B compounds [56], multiple macrolide-resistance genes were identified. The most abundant were *ermB*, (835/1271; 66%), *ermT* (14.7%), and *ermA* (6.7%) (Figure S3). Some of these genes showed a bias for the CLIN (ermT) or AGRI and WW-AGR (ermA) environments, while *ermB* was prevalent in all environments except NWS and WW-MUN from AB. The majority of these genes were localized on plasmids, with *ermA* identified on AB369 (44/47;94%) *ermB* mostly associated with AC731 (146/662; 22%) and AB369 (130/662; 20%). In AGRI genomes, *ermB* was commonly associated with AC730 (37/76; 49%) and AB756 (15/76; 20%). Plasmid clusters AC731 and AB369 were also associated with *vanA* and *ermB*. The *ermA* gene, was exclusively associated with Type A and the AGRI isolates, while *ermB* was positively associated with Type A, as well as with UK, AGRI, NWS, CLIN and WW-MUN isolates. The *ermT* gene demonstrated a positive association with Type A, UK, and CLIN isolates and a negative association with AGRI and NWS isolates.

**Fig. 6.**
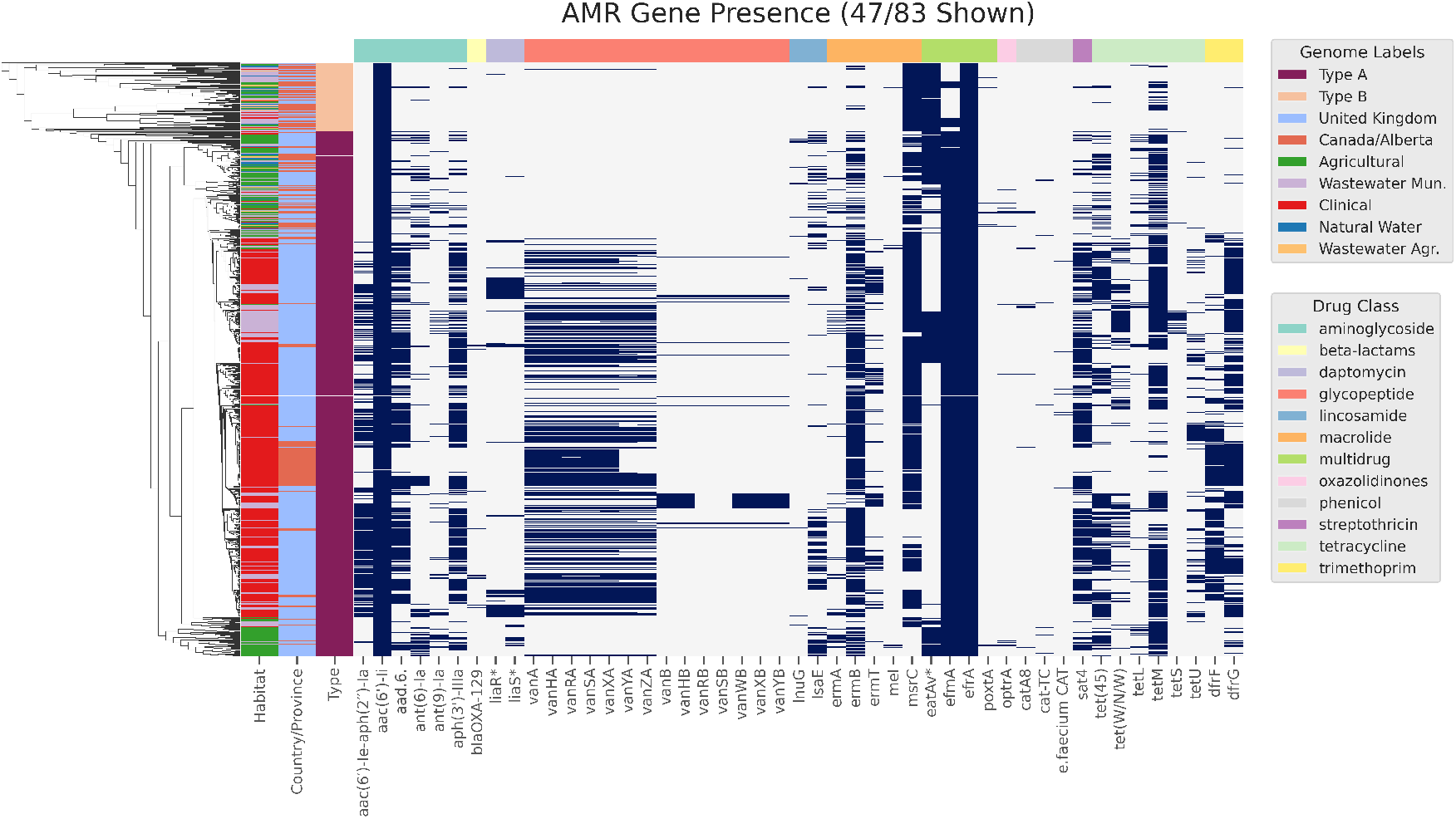
Heatmap showing the presence of AMR determinant genes detected in 1273 *E. faecium* genomes analyzed in this study. The y-axis indicates genomes (color coded by habitat, geography and type) sorted by topology of the core genome maximum likelihood tree. AMR determinants (x-axis) are sorted by drug class. * denotes variant versions of intrinsic genes conferring AMR.

Tetracycline-resistance genes were common, often plasmid-associated, in the analyzed genomes (Figure S4). *tetM* (783/1273; 61.5%) was most prevalent, followed by *tet(45)* (455/1273; 35.7%) in CLIN genomes from the UK. Other tetracycline genes including *tet(W/N/W)* (236/1273; 18.5%) and *tetU* (168/1273; 13.2%) were found at higher rates in WW-MUN and CLIN genomes from the UK; *tetL* (90/1273; 7.1%) was found mostly in agriculture and WW-AGR with higher levels in AGRI genomes in the UK; *tetS* (37/1273; 2.9%) was found primarily in WW-MUN and NWS. *tet(40)* and *tetO* were both found in UK WW-MUN.

The aminoglycoside resistance gene *aac6-Ii*, responsible for intrinsic aminoglycoside resistance in this species [57], was found in the majority of genomes (1271/1273; 99.8%) (Figure 6). Interestingly, a three-gene locus *aad(6)-sat4-aph(3’)-IIIa* (595/1273; 46.7%) conferring resistance to aminoglycosides and streptothricin, was present in AGRI, CLIN, and WW-MUN isolates at higher prevalence in UK than AB. A bi-functional protein-coding gene *aac(6’)-Ie-aph(2”)-Ia* (468/1273; 36.8%) was also found at higher abundance in the UK CLIN and WW-MUN isolates compared to AB. Both *ant(6)-Ia* (166/1273; 13.0%) and *ant(9)-Ia* (82/1273; 6.4%) exhibited a higher prevalence in AGRI isolates than isolates from other habitats. A small number of CLIN and WW-MUN isolates from UK harboured *ant(9)-Ia*, while corresponding Alberta isolates lacked this gene. Other rarely detected aminoglycoside-resistance genes included *apmA* (11/156 UK AGRI genomes), *aph2-IVa* (2/270 UK WW-MUN genomes), *aac(6)-Iak* (2/544 UK CLIN and 2/270 UK WW-MUN genomes), *aac(6)-II* (1/544 UK CLIN genomes), *aph2-Ie* (1/270 UK WW-MUN genomes), and *ant(4)-Ib* (2/51 AB AGRI genomes).

### 3.8 Heavy Metal and Biocide Resistance Genes

Genes related to copper resistance and transport comprised a large portion of the predicted HMR genes comprising 12/32 gene clusters and 2542/7914 predicted genes. The most common were two *copB* clusters (BacMet clusters 3 and 9) and a *copA* cluster (BacMet cluster 5). While cluster 3 *copB* were spread across habitats and countries, the cluster 9 *copB* were most predominant in AB WW-MUN, AGRI, NWS, and WW-AGR isolates and were underrepresented in CLIN and all UK isolates (Figure S5). The *copB* genes chromsomal except for 2/1155 cluster 3 and 3/137 cluster 9 *copB* representatives associated with GIs and plasmids, respectively. The *copA* cluster 5 was also prevalent in all environments, although more so in CLIN isolates and UK WW-MUN. Most *copA* (549/917) were also predicted to be associated with the chromosome while 366/917 cases were predicted to be localized to plasmid AH273. Another cluster of copper-resistance genes primarily associated with the agricultural environment in the UK included the genes *mco* (BacMet cluster 14), *tcrB* (BacMet cluster 15), *tcrA* (BacMet cluster 16), and *copY/tcrY* (BacMet cluster 17). These genes all showed strong association with one another as well as the plasmid AC726. There were five instances identified where copper genes and mercury-resistance genes were colocalized on a single MGE, with plasmid cluster AD908 involved in three instances. All of these cases were identified in AGRI genomes from the UK.

### 3.9 Virulence Factors

Both the *ssaB* and *fss3* genes have been shown to play a role in adhesion. These genes were primarily identified in CLIN genomes from both countries and WW-MUN genomes from the UK. *ssaB* was either localized on the chromosome (364/627; 42%) or on GIs (263/627; 58%) (Figure S6). In particular, 59% (215/263) of the genes were associated with a single GI. An additional 36% (95/263) were identified on GI 23, which also carried *fss3* in 85% (81/95) of cases. GI 23 was primarily identified in the UK dataset (93/95). *fss3* was found in 63% of UK CLIN genomes but only 14% of AB CLIN genomes. Among the remaining *fss3* genes not present on GI 23, 37% (181/483) were present on the chromosome, 27% (132/483) on other GIs, 18% (88/483) on regions predicted to be both a GI and a prophage, and one predicted to be on a non-GI-associated prophage. Both *ssaB* and *fss3* were strongly correlated to each other, clinical-related AMR genes, and MGEs.

A total of 25 VFDB gene clusters were predicted to be pilin genes common to all habitats. The CLIN genomes had the highest prevalence of these genes and the proportion of UK AGRI isolates with each of these genes was higher than the AB AGRI isolates (Figure S7). Plasmids were the most common localization site of *pilA* (929/958; 97.0%), *pilE* (1023/1060; 96.5%), and *pilF* (881/909; 96.9%), with the most common colocalized plasmids being AD907 and AC731. The chromosomal *pilB* gene had a similar prevalence across all datasets and was found on the chromosome.

## 4 Discussion

*E. faecium*, commonly a minor harmless component of the enteric microbiota, has become a leading causative agent of healthcare-associated infections since the early 1980s [58, 59]. The combination of approaches we applied here can augment phenotype-based “One Health” genomic-surveillance workflows for *E. faecium* and other bacterial pathogens. Using the whole-genome approach the potential for gene transfer and the distributions of genes within and between habitats can be defined.

### 4.1 Genomic Epidemiology Suggests Strong Habitat Associations But Few Barriers to Transmission

The AB and UK genomes are distributed on the core genome-based phylogenetic tree independent of type or habitat. This indicates that geographic separation has very little impact on the population structure of *E. faecium*. There was no significant difference in the abundance of MGEs between types or geographic origin (Table 1, 2) which appears to contradict the findings of previous studies that detected significantly higher numbers of MGEs in Clade A than in Clade B [21, 27, 60]. The phylogenetic approach of BayesTraits suggested that the observed differences were driven by increased MGE abundance in CLIN and WW-MUN isolates (Figure 3). The lack of observed association with type suggests that MGEs can move between phylogenetically distant *E. faecium* isolates and that MGEs within populations are similar across continental barriers.

The global patterns of association seen among AMR and VF genes mirrored those of MGEs, with habitat generating the smallest p-values in the ANOVA tests (Tables 1 and 2). Geographic origin showed strong associations only in interactions with habitat for AMR genes and MGEs, likely as a result of intensive use of antimicrobials which can result in the emergence of multidrug-resistant *E. faecium*. *E. faecium* may acquire a multi-drug resistance plasmid in the clinic but lose it upon introduction to another habitat [61]. This process can occur rapidly and repeatedly, with habitat serving only as an ecological filter rather than a barrier to transmission. Analysis of the composition of features by groupings of type, habitat, and geography supports the division of features based on type and habitat.

Antimicrobial use may account for some of the variance in the distribution of AMR genes. The few differences we observe may be due to different host origins in the two locations: the UK AGRI genomes include isolates from chicken, turkey, pig, beef, and dairy cattle while the AB AGRI samples only originated from beef cattle production sources. Second, antimicrobial use in agriculture differs between Alberta and the UK. For example, the strongest associations of AMR genes with location were among the aminoglycoside family (e.g., *aad(6*)), which are used in the UK but not Alberta [23, 24]. However, our information about antibiotic use is incomplete, especially for chicken and turkey [62, 63].

### 4.2 A Highly Diversified and Dynamic Accessory Genome Allows the Rapid Acquisition of Resistance and Other Traits

Although some of the 88 plasmid clusters and 824 GI sets identified are likely very similar, they are nonetheless different enough in sequence and gene content to be differentiated. The number of unique prophages is likely an underestimate due to the grouping of some phages by name (e.g., *‘Streptococcus* phage’) rather than by homology. Key resistance genes were observed in association with many predicted plasmid clusters. For example, 14 observed clusters had at least one *vanA* gene. Similarly, multiple clusters showed associations with tetracycline, aminoglycoside, and macrolide-resistance genes. The dispersion of genes across multiple clusters likely diminishes the strength of observed associations, such that we may not identify all MGEs that associate with specific genes. Additionally, aggregation of plasmids and GIs into broader clusters (such as plasmid incompatibility groups) and consequently fewer classes might improve our ability to detect important associations. The increased prevalence of many classes of resistance genes and MGEs in the clinical environment suggests that this may be the key focal point of plasmid evolution, with novel combinations forming through recombination events. The lack of geographic barriers, and the apparent ability of *E. faecium* to move between habitats, suggests that new MGEs will not be limited in their ability to disperse. The clear ability of *E. faecium* to acquire and disseminate new genes from distantly related species creates additional risks for the emergence of new combinations of AMR determinants.

### 4.3 An Emerging Clade of Pathogenic E. *faecium*

The groEL-based clade-mapping on our reference tree supports a monophyletic Clade A, as described and proposed by Palmer *et al*. [64], but a paraphyletic “Clade B”, which has led to our designation of these two groups as “types” rather than clades. Earlier work proposed a division of Clade A into a pathogenic subclade A1, and a commensal group specific to non-human animals capable of causing sporadic infections, subclade A2 [21]. However, consistent with recent observations by others, we observed a large group comprised of CLIN and WW-MUN isolates that branched within the larger grouping that included genomes isolated from all habitats (Type A); this tree topology has been referred to as a clonal expansion [65, 66]. Importantly, our phylogenetic tree is based on the core genome of our isolates (n=1273) and, therefore, its topology should be less affected by LGT events than a gene-focused or whole-genome tree and thus should better reflect the structure and evolutionary trajectory of *E. faecium* populations.

Our core-genome tree suggests that there may be a sub-population adapting to the clinical niche beyond the simple and more plastic advantage provided by MGEs. This is consistent with the observation from Leclerque *et al*. that members of this clinical expansion are out-competed by other *E. faecium* clones in natural environments [67] and by Montealegre *et al*. that strains from Type B have higher fitness than Type A in the absence of antibiotics [68]. These findings, together with our results, support the hypothesis proposed by Prieto *et al*. that this clinical clonal expansion is so specialized to its environment that its strains are unfit to populate other environments and niches. This population could get more isolated and drift away from the rest of the species [66]. If the clinical-associated group we detected in our dataset has some level of diversification, it may satisfy the Cohan & Perry (2007) definition of an ecotype, with lineage cohesiveness conferred by genetic similarities and distinguished by unique adaptations (e.g., AMR genes) and ecological capabilities. Cohan & Perry hypothesized that periodic selective sweeps reduce genetic variation between the genomes of organisms specialized to certain ecological niches, increasing the differences between these ecotypes and the rest of the named species [69]. However, other authors emphasize the role of recombination in bacterial divergence which allows for gene-specific sweeps [70].

Other pathogenic species can give some insight into the driving forces shaping the *E. faecium* population structure we observed. For example, in the last 30 years a new multidrug resistant genotype, H58, of *Salmonella enterica* sp. *enterica* sv. Typhi (*S*. Typhi) emerged as a clonal expansion and has rapidly spread globally [71]. This clade, like *E. faecium*, has rapidly differentiated into major antimicrobial-resistant lineages [72], but, unlike *E. faecium*, the differentiation has been driven by strong geographical selection [71]. Interestingly and contrary to *E. faecium*, H58 has a similar fitness to other *Salmonella* genotypes in absence of antimicrobials and local genetic drift rather than niche specialization is responsible for its diversification. This difference suggests that the *E. faecium* clinical ecotype is locked into clinical associated environments as competitive exclusion from other strains prevent its expansion [67, 68]. However, relying solely on this niche restriction to guide our surveillance efforts would be a mistake, as recombinatory events are common in *E. faecium* [73] and several circulating non-clinical Type A strains are likely recombinants between Type A and Type B [74]. Therefore, it is possible for neglected non-pathogenic strains to acquire the traits necessary to expand to the clinical environment while retaining their cosmopolitan lifestyle.

### 4.4 Towards Monitoring of Evolving Threats

As in many other pathogens, the genomic plasticity of *E. faecium* effectively creates a reservoir of genes and MGEs that can reshuffle according to environmental pressures and opportunities, with geographic distance, phylogenetic distance, and habitat boundaries as no obstacle. Although the analytical pipeline we apply here was effective in detecting environmental and genetic connections, improvements in sampling, sequencing, and analysis will be needed to realize the full potential of genomic monitoring. While it makes sense to focus efforts on the sampling of clinical isolates, isolates from other environments need to be collected with appropriate metadata such as local antimicrobial usage and connectivity patterns with other sampling locations.

The limitations of short-read sequencing are well documented, and MGEs are generally more difficult to recover due to the increased abundance of repeat regions [75–77]. Hybrid long-read / short-read assemblies can provide complete or near-complete information about MGE gene content, and enrich reference databases to serve as references for short-read assemblies. Future work should also include refinements of the statistical methods used and techniques to identify key genes. For example, contextual information such as gene order can enhance the differentiation of true AMR genes from highly similar false positives. In our analysis, we found that the filtering parameters applied to the analytical outputs are of paramount importance. In fact, after curation, some of the more rare genes we identified proved to be artifacts generated by thresholds that were not stringent enough to root out low levels of sequence contamination or short but unreliable matches to databases.

## 5 Conclusion

Our examination of the evolution of *E. faecium* revealed a great capacity to acquire traits such as antimicrobial resistance, fueled by a large repertoire of MGEs. Misuse of antimicrobials has driven *E. faecium*’s transformation to a major contributor to morbidity and mortality worldwide. The datasets we consider here, when combined with other large collections, provide a robust snapshot of the current population structure and evolutionary trends of *E. faecium* across the One Health continuum providing crucial insight to target surveillance, design public health policies, and inform interventions. The work we present here can support the careful stewardship, comprehensive monitoring, and measurable outcomes that will be necessary to manage *E. faecium* and other evolving pathogens.

## 6 Author statements

### 6.1 Authors and contributions

H.S., K.G., F.S.L.B., R.C.F., R.Z., and R.G.B. conceptualized the study. H.S., K.G., A.M., F.M., A.K., C.L., C.N.R., J.H.E.N., J.R., K.B., M.O., B.P.A., A.R.R., A.G.M., F.S.L.B., R.C.F., and R.G.B. contributed one or more of experimental design and execution, development and validation of software, and data analysis. H.S., F.M., T.A.M., S.J.P., K.E.R., T.G., and R.G.B. contributed datasets and/or performed curation of datasets. H.S., K.G., A.M., A.K., R.C.F., R.Z., and R.G.B. prepared the initial draft of the manuscript. All authors edited and approved the final version of the manuscript.

### 6.4 Data summary

The code and data to reproduce the results are available at GitHub (https://github.com/beiko-lab/efaecium-niche). Supplementary Data are available at [WHERE]

## 6.5 Acknowledgements

We thank Henry and Charlie Sanderson for their constant support.

