## Supplementary Figures for "Exploring the mobilome and resistome of *Enterococcus faecium* in a One Health context across two continents"

S2 BayesTraits *p* value density plots, panels A-B . . 5
S2 BayesTraits *p* value density plots, panels C-D . . 6
S2 BayesTraits *p* value density plots, panels E-F . . 7
S3 Macrolide resistance detailed plots. . . . . . . . . . . 8
S4 *tet* gene detailed plots. . . . . . . . . . . . . . . . . . . . . . 9
S5 *cop* gene detailed plots. . . . . . . . . . . . . . . . . . . . . 10
S6 Virulence factor genes detailed plots. . . . . . . . . . 10
S7 *pil* gene detailed plots. . . . . . . . . . . . . . . . . . . . . . 11

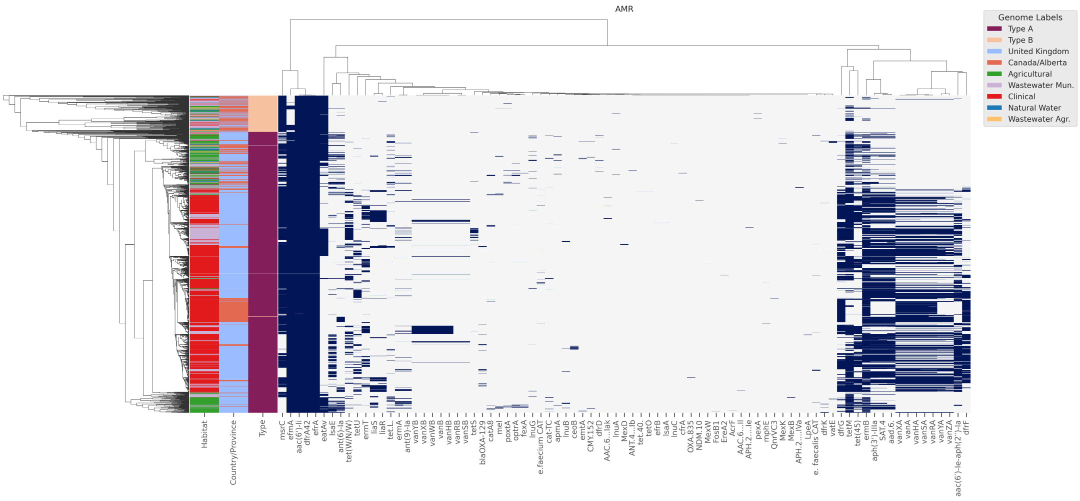

(a) AMR determinants.

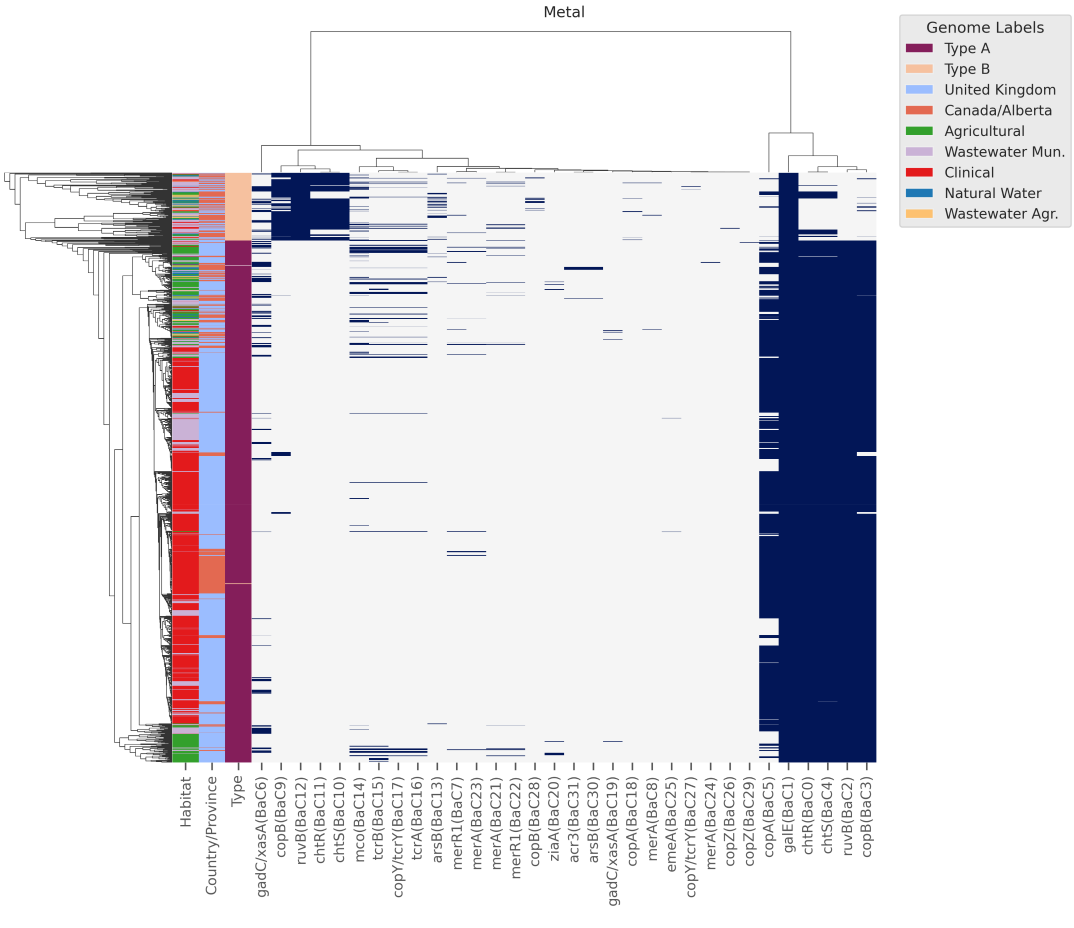

(b) HMR Determinants.

**Figure S1.** Feature clustergrams of target genes and MGEs. Rows indicate genomes, sorted by topology of the core genome maximum likelihood phylogenetic tree. Row colors indicate genome habitats, country of origin, and type assignment from left to right. Columns are clustered using single-linkage agglomerative clustering with Manhattan distance over the presence/absence vectors. (A) AMR determinants. (B) HMR determinants. (C) Virulence factors. (D) Plasmid clusters. (E) Genomic islands. (F) Phages.

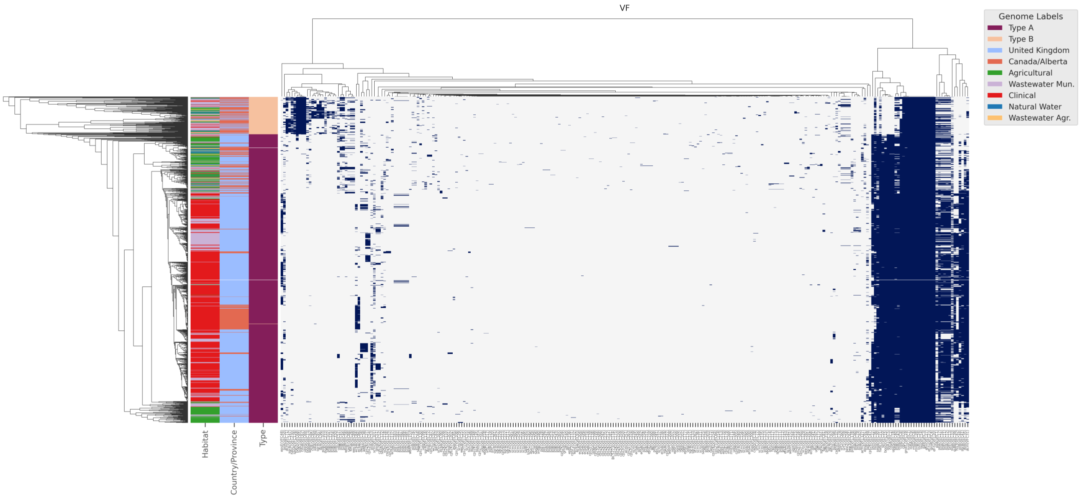

(c) Virulence factors.

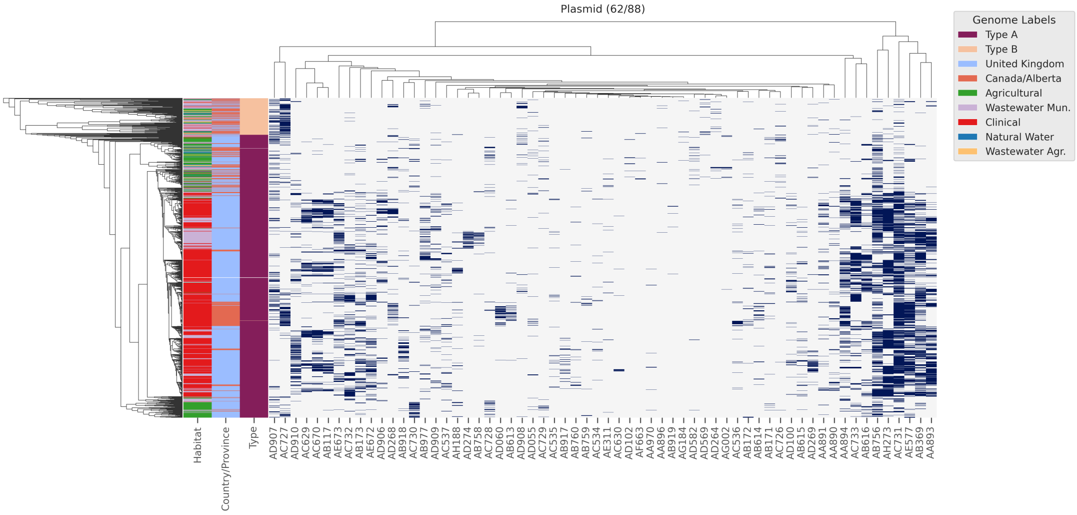

(d) Plasmid clusters.

**Figure S1.** Feature clustergrams of target genes and MGEs. Rows indicate genomes, sorted by topology of the core genome maximum likelihood phylogenetic tree. Row colors indicate genome habitats, country of origin, and type assignment from left to right. Columns are clustered using single-linkage agglomerative clustering with Manhattan distance over the presence/absence vectors. (A) AMR determinants. (B) HMR determinants. (C) Virulence factors. (D) Plasmid clusters. (E) Genomic islands. (F) Phages.

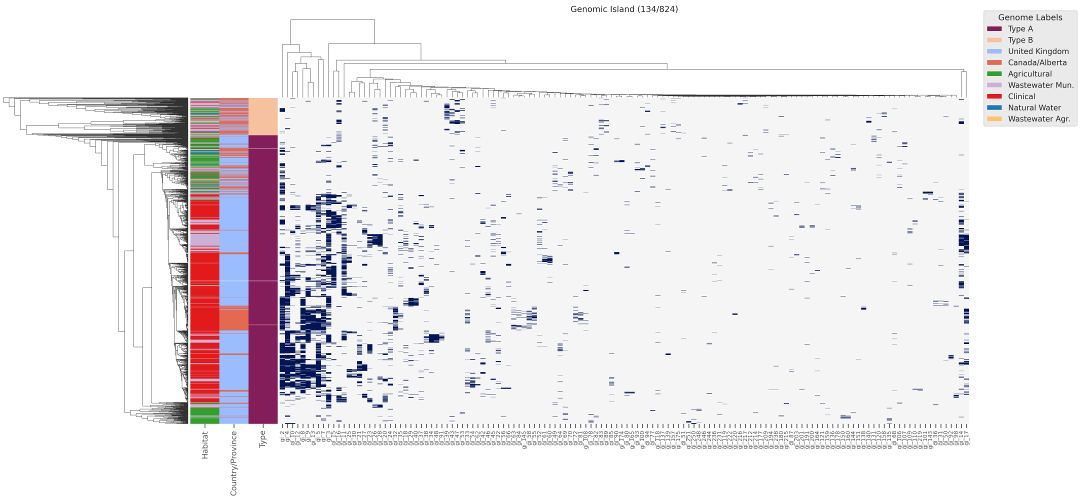

(e) Genomic Islands.

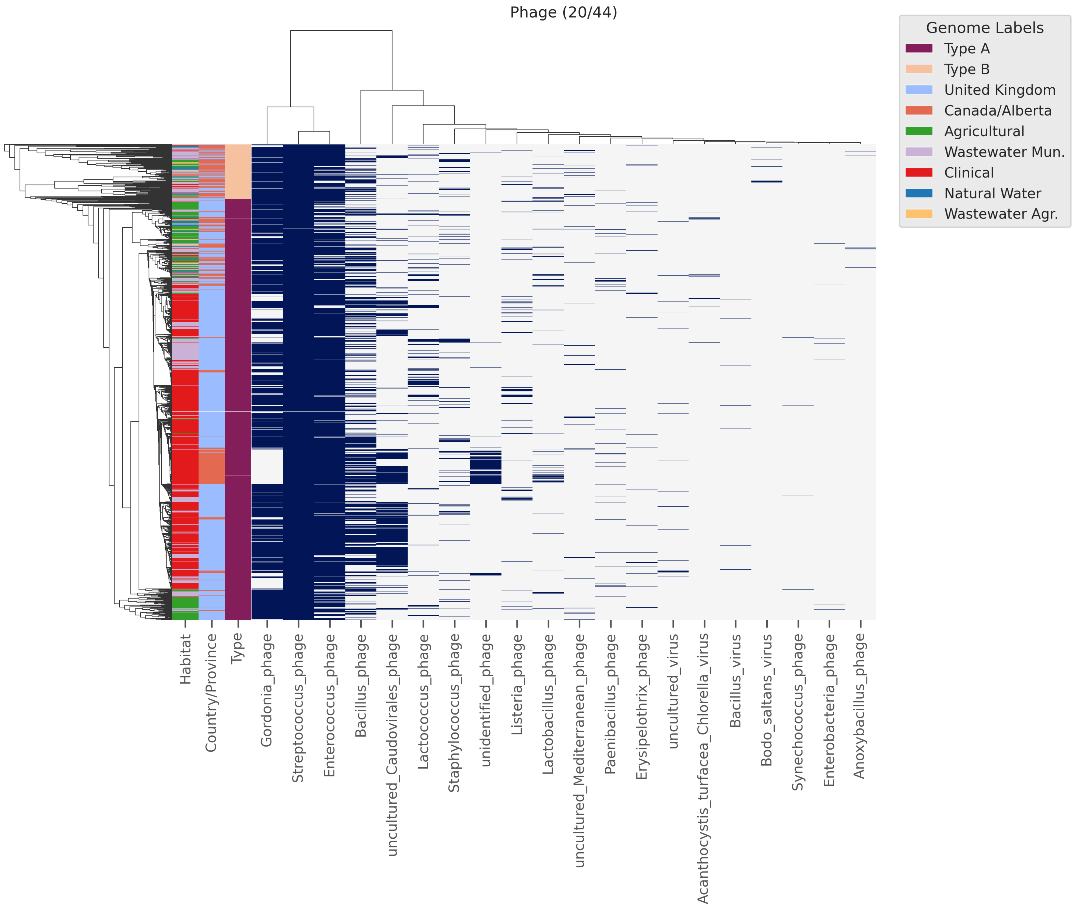

(f) Phages.

**Figure S1.** Feature clustergrams of target genes and MGEs. Rows indicate genomes, sorted by topology of the core genome maximum likelihood phylogenetic tree. Row colors indicate genome habitats, country of origin, and type assignment from left to right. Columns are clustered using single-linkage agglomerative clustering with Manhattan distance over the presence/absence vectors. (A) AMR determinants. (B) HMR determinants. (C) Virulence factors. (D) Plasmid clusters. (E) Genomic islands. (F) Phages.

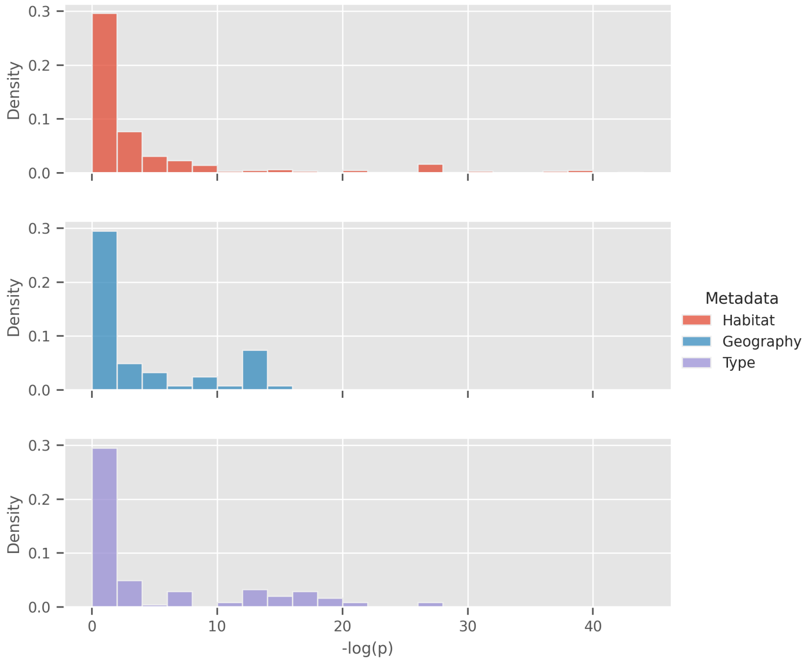

(a) AMR determinant $p$ value distributions.

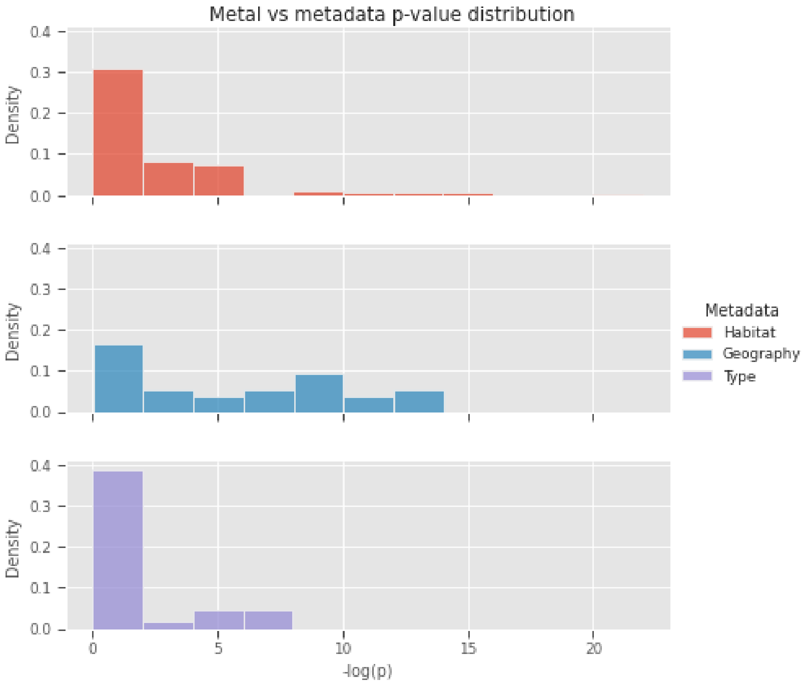

(b) HMR determinant $p$ value distributions.

**Figure S2.** Density histograms of BayesTraits $p$-values testing the hypothesis that feature evolution is dependant on habitat category, geographical sampling location, and assigned type. Y axis indicates the total are under the curve for each histogram. X axis indicates $-log(p)$. (A) AMR determinant genes. (B) HMR determinant genes. (C) Virulence factors. (D) Plasmid cluters. (E) Genomic islands. (F) Phages.

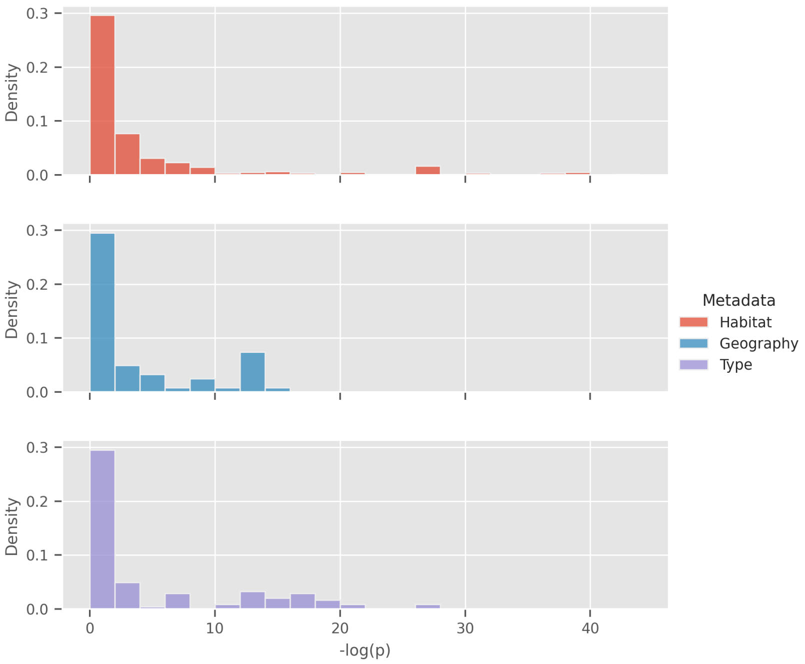

(c) Virulence factor $p$ value distributions.

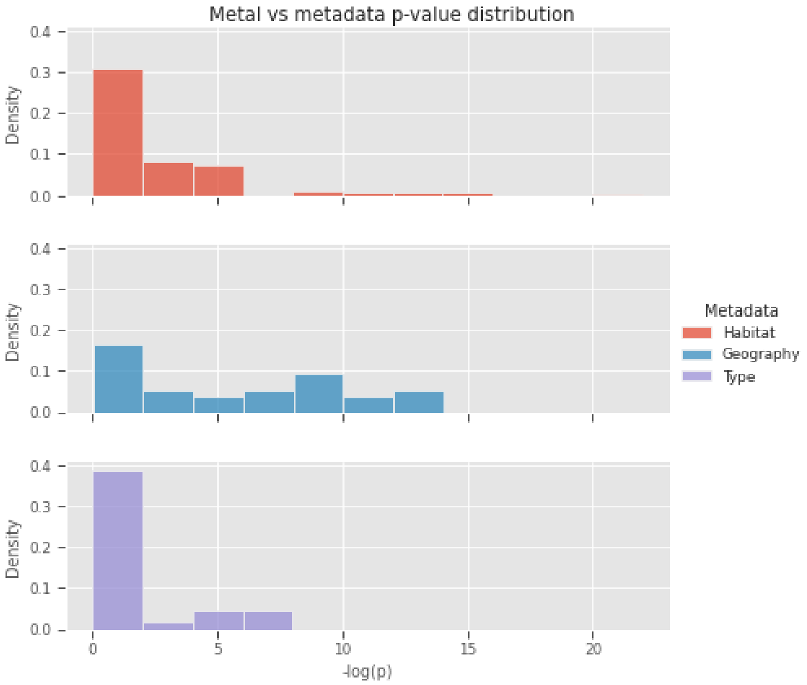

(d) Plasmid cluster $p$ value distributions.

**Figure S2.** Density histograms of BayesTraits $p$-values testing the hypothesis that feature evolution is dependant on habitat category, geographical sampling location, and assigned type. Y axis indicates the total are under the curve for each histogram. X axis indicates $-log(p)$. (A) AMR determinant genes. (B) HMR determinant genes. (C) Virulence factors. (D) Plasmid cluters. (E) Genomic islands. (F) Phages.

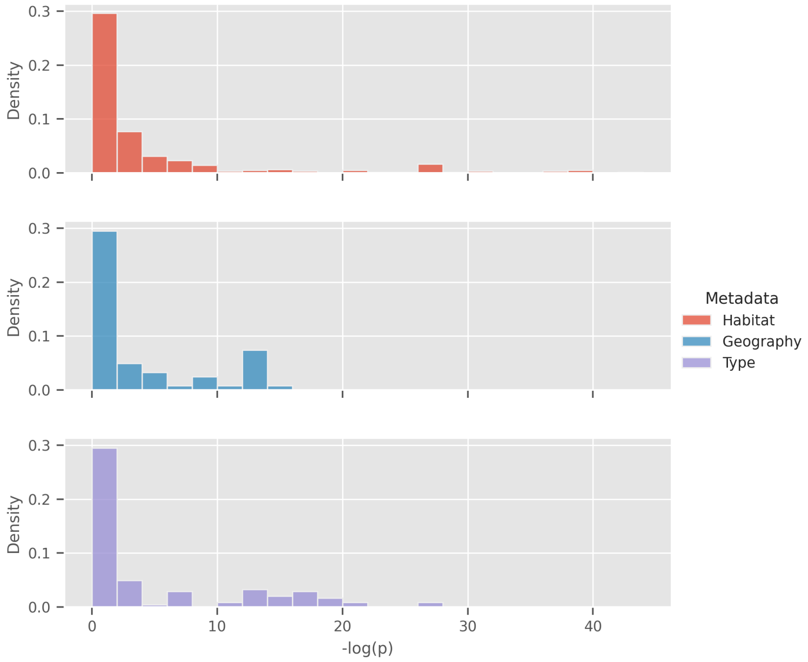

(e) Genomic island $p$ value distributions.

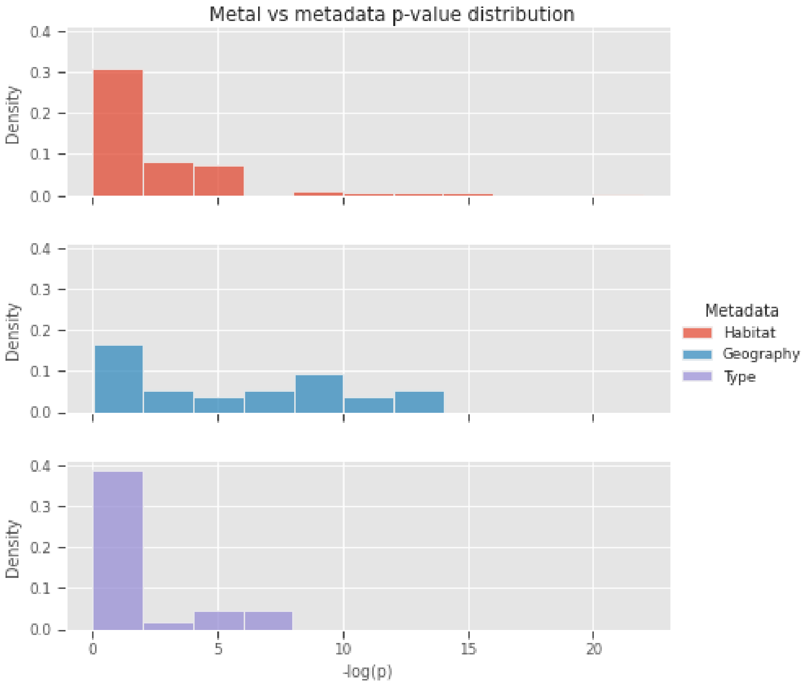

(f) Phage $p$ value distributions.

**Figure S2.** Density histograms of BayesTraits $p$-values testing the hypothesis that feature evolution is dependant on habitat category, geographical sampling location, and assigned type. Y axis indicates the total are under the curve for each histogram. X axis indicates $-log(p)$. (A) AMR determinant genes. (B) HMR determinant genes. (C) Virulence factors. (D) Plasmid cluters. (E) Genomic islands. (F) Phages.

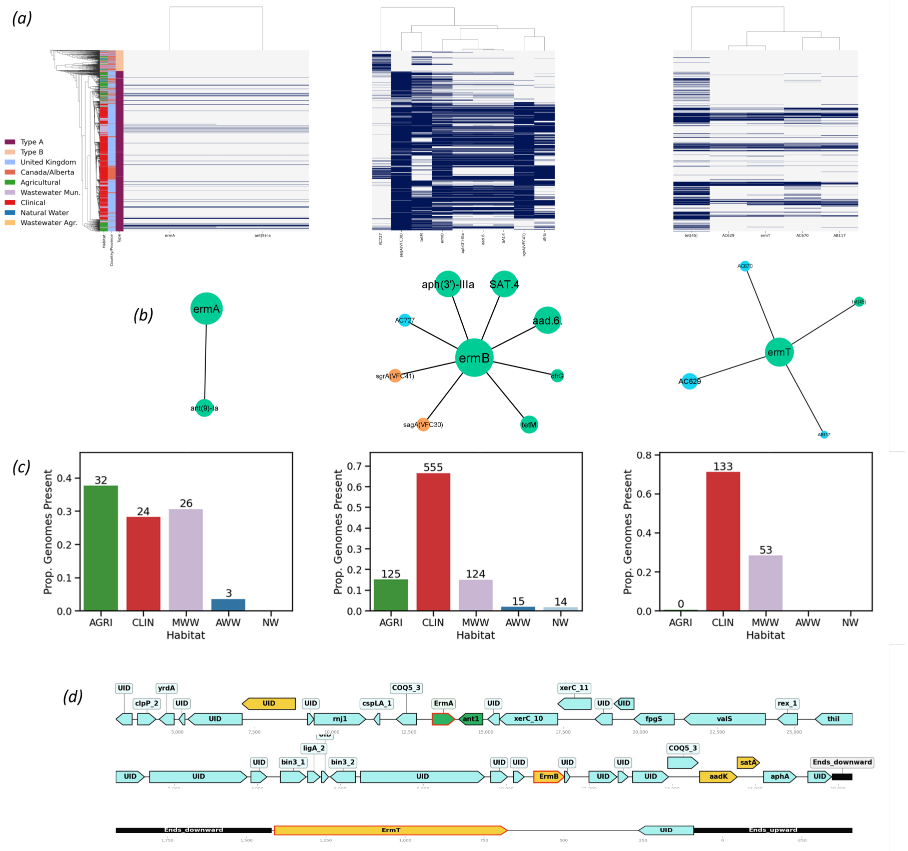

**Figure S3.** Statistical associations and physical localization of macrolide-resistance genes *ermA*, *ermB*, and *ermT*. (a) phylogenetic distribution of erm genes and associated genes with associated likelihood ratio $\geq$ 100. (b) Statistical association networks of *erm* genes. (c) Distribution of genes by habitat. (d) Examples of gene order on identified contigs. Gene and MGE colours are consistent with those in Figure 2. Green genes correspond to perfect matches with reference genes in the CARD database, yellow genes are Strict hits.

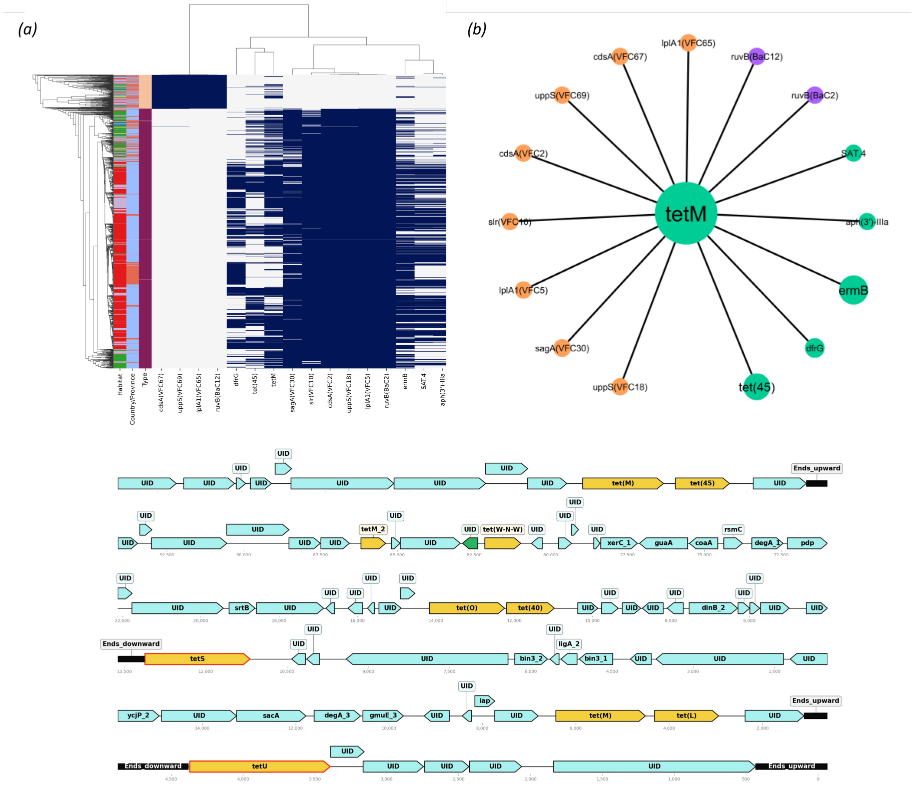

**Figure S4.** Associations of tetM with other features and example neighbourhoods of tet genes. (a) phylogenetic distribution of tetM and genes with associated likelihood ratio $\geq$ 100. (b) Statistical association networks of tetM. (c) Co-localization of tetM with other tetracycline-resistance genes, and example neighbourhoods of other tet genes. Gene and MGE colours are consistent with those in Figure 2. Green genes correspond to perfect matches with reference genes in the CARD database, yellow genes are Strict hits.

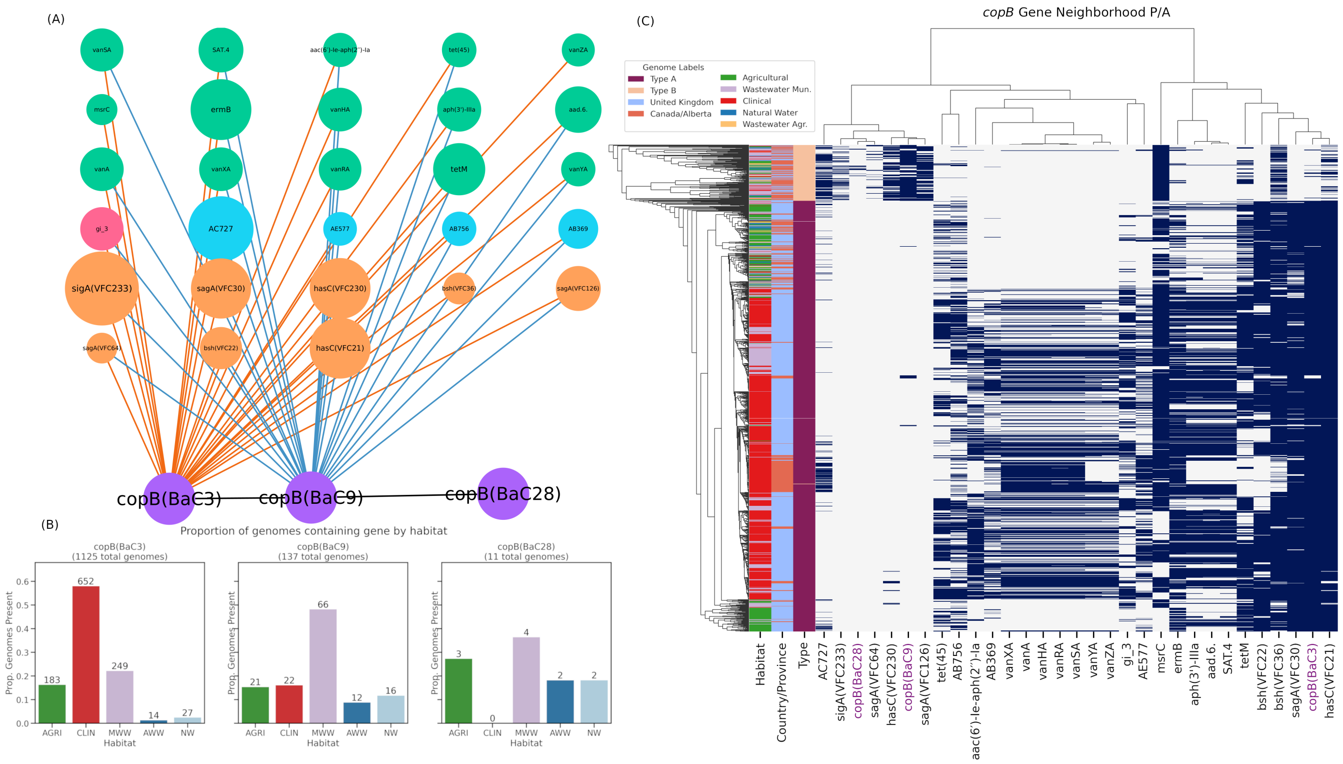

**Figure S5.** Distribution and associations of *copB* genes. (A) Statistical association network of *copB* genes with other features. Gene and MGE colours are consistent with those in Figure 2. (B) Habitat distributions of *copB* genes. (C) Phylogenetic distribution of genes and other features with an associated likelihood ratio $\geq 50$.

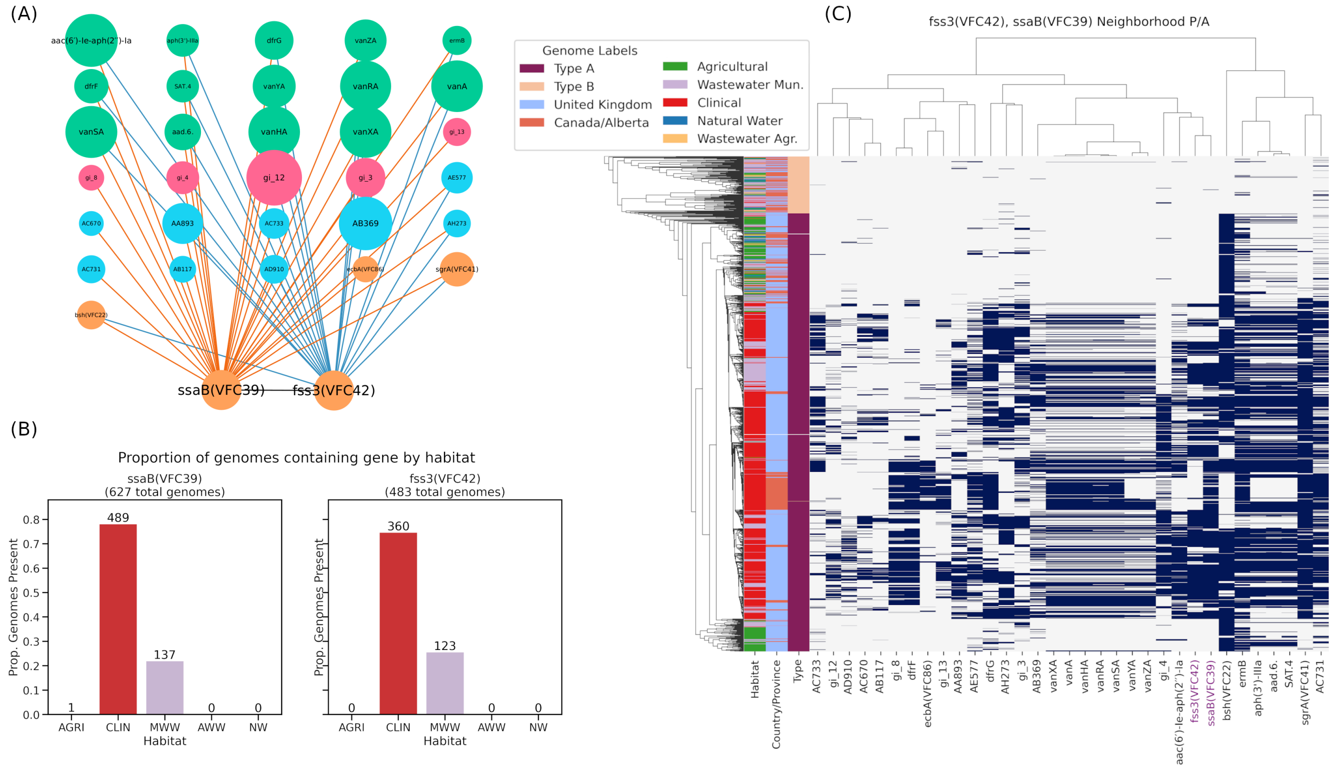

**Figure S6.** Distribution and associations of *ssaB* and *fss3* genes. (A) Statistical association network of *ssaB* and *fss3* genes with other features. Gene and MGE colours are consistent with those in Figure 2. (B) Habitat distributions of *ssaB* and *fss3* genes. (C) Phylogenetic distribution of genes and other features with an associated likelihood ratio $\geq 100$.

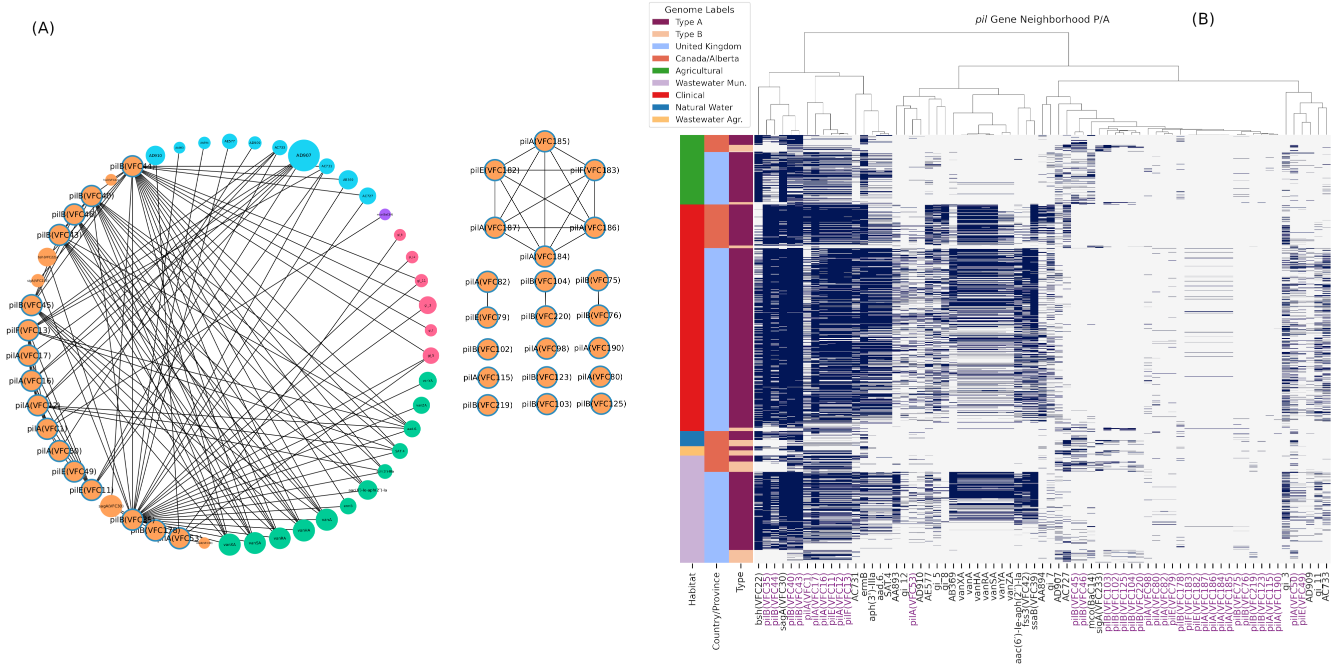

**Figure S7.** Distribution and associations of *pil* genes. (A) Statistical association network of *pil* genes with other features. *pil* genes are circled in blue. Gene and MGE fill colours are consistent with those in Figure 2. (B) Presence/absence of *pil* genes. Rows indicate genomes and are sorted by habitat. Genes include *pil* genes and genes/MGEs that associate with any *pil* gene with a likelihood ratio $\geq 75$.
